# CytoPy: an autonomous cytometry analysis framework

**DOI:** 10.1101/2020.04.08.031898

**Authors:** Ross J. Burton, Raya Ahmed, Simone M. Cuff, Sarah Baker, Andreas Artemiou, Matthias Eberl

**Author notes:** Correspondence to: Ross Burton, Division of Infection and Immunity, Henry Wellcome Building, School of Medicine, Cardiff University, Heath Park, Cardiff CF14 4XN, Wales, United Kingdom.

## Abstract

Cytometry analysis has seen a considerable expansion in recent years in the maximum number of parameters that can be acquired in a single experiment. In response to this technological advance there has been an increased effort to develop new computational methodologies for handling high-dimensional single cell data acquired by flow or mass cytometry. Despite the success of numerous algorithms and published packages to replicate and outperform traditional manual analysis, widespread adoption of these techniques has yet to be realised in the field of immunology. Here we present CytoPy, a Python framework for automated analysis of cytometry data that integrates a document-based database for a data-centric and iterative analytical environment. In addition, our algorithm agnostic design provides a platform for open-source cytometry bioinformatics in the Python ecosystem. We demonstrate the ability of CytoPy to phenotype T cell subsets in whole blood samples even in the presence of significant batch effects due to technical and user variation. The complete analytical pipeline was then used to immunophenotype the local inflammatory infiltrate in individuals with and without acute bacterial infection. CytoPy is open-source and licensed under the MIT license. CytoPy is open source and available at https://github.com/burtonrj/CytoPy, with notebooks accompanying this manuscript (https://github.com/burtonrj/CytoPyManuscript) and software documentation at https://cytopy.readthedocs.io/.

## 1. Introduction

Cytometry data analysis has undergone a paradigm shift in response to the growing number of parameters that can be observed in any one experiment. As the field evolves, the traditional method of manual gating by sub-setting single cell data into populations and encircling data points in hand-drawn polygons in two-dimensional space is proving laborious, subjective, and difficult to standardise. In response to these shortcomings, a cross-disciplinary effort has given birth to a new approach often termed ‘cytometry bioinformatics’, to leverage complex computer algorithms and machine learning to automate analysis and improve the investigator’s ability to extract meaning from high dimensional data.

Where cytometry is used for data acquisition, the typical objective is to discern differences between groups of subjects or experimental conditions, or to identify a phenotype that correlates with an experimental or clinical endpoint. To this end, a computational approach to analysis of cytometry data can take one of two strategies: to group events based on similarity (*e.g.* cell populations), which then form the variables (often descriptive statistics of the obtained groups) the investigator uses to test their hypothesis, or directly model the acquired multi-dimension distribution with respect to a chosen endpoint. Classification strategies can be further subdivided: autonomous gating replicates traditional gating by applying algorithms to data in one or two dimensions (flowDensity [1], OpenCyto [2]); clustering in high-dimensional space to group events according to their individual characteristics (FlowSOM [3], Phenograph [4], Xshift [5], SPADE [6]); and supervised or semi-supervised classification where manual annotations are used to train a model capable of identifying cell populations within unlabelled data (FlowLearn [7], ACDC [8], DeepCyTof [9]). Direct modelling strategies have been successfully adopted in applications such as ACCENSE [10], CellCNN [11], CytoDX [12] and in the work described by Hu *et al.* [13]. This approach has the benefit of removing any subjectivity and can be considered as truly automated but requires the pooling of sample data and is therefore sensitive to batch effects.

In addition, various pieces of software and pipelines have been developed for data handling, transformation, normalisation and cleaning (*e.g*. flowCore, flowIO, flowUtils, flowTrans, reFlow, flowAI), visualisation (*e.g*. ggCyto, t-SNE, UMAP, PHATE), and specific applications (*e.g.* Citrus, MetaCyto, flowType/RchyOptimyx) [14]. However, widespread adoption of cytometry bioinformatics has yet to be realised and a lack of consensus remains on how to implement such technologies across the scientific community, with much of the analysis pipeline left to the individual investigator to establish. This inconsistency continues to result in projects amassing collections of custom scripts and data management that are not standardised or centralised, which not only makes reproducing results difficult but also makes for a daunting landscape for newcomers to the field.

We here introduce ‘CytoPy’, a novel analysis framework that aims to mend these issues whilst granting access to state-of-the-art machine learning algorithms and techniques widely adopted in cytometry bioinformatics. CytoPy was developed in the Python programming language, which prides itself on readability and a beginner friendly syntax. CytoPy introduces a central data source for all single cell data, experimental metadata, and analysis results, and provides a ‘low code’ interface that is both powerful and easy to maintain. CytoPy incorporates popular data science and machine learning libraries such as Pandas [15], Scikit-Learn [16] and Tensorflow [17], with an application programming interface (API) designed to help expand cytometry bioinformatics in the Python ecosystem. In addition, CytoPy provides convenient access to popular algorithms and techniques popular in cytometry data analysis such as PHATE [18], UMAP [19], Phenograph [4], and FlowSOM [3].

A burgeoning challenge as cytometry data grows in size is batch effect. Nowhere is this issue more pressing than in translational research where lengthy study designs and complex specimens make technical variation unavoidable. CytoPy provides tools for visualising and quantifying batch effect as well as methods to subvert and eliminate it. To validate CytoPy we used in-house data obtained from patients undergoing peritoneal dialysis and who presented with and without acute bacterial infection. Data had been collected over several years, therefore providing ample opportunity to demonstrate the ability of CytoPy to navigate issues such as batch effects and inconsistent meta-data. We first introduce the capabilities of CytoPy by characterising T cells in blood and comparing findings to manual gates. Finally, we employ the entire pipeline to describe a known phenotype of immune cells in the peritoneal effluent of dialysis patients that differentiates individuals with acute peritonitis from stable controls. Accompanying Jupyter Notebooks demonstrating all necessary code are available here at https://github.com/burtonrj/CytoPy_Peritonitis. We believe that CytoPy provides a powerful and user-friendly framework to interrogate high dimensional data originating from investigations using flow cytometry or mass cytometry as readout, and has the potential to facilitate automated data analysis in a multitude of experimental and clinical contexts.

## 2. Design and Implementation

### 2.1 Building a framework that is algorithm-agnostic and data-centric

Reliable data management is a cornerstone of successful analysis, improving reproducibility and fostering collaboration. A typical cytometry project consists of many Flow Cytometry Standard (FCS) files, clinical or experimental metadata, and additional information generated throughout the analysis (*e.g*. gating, clustering results, cell classification, sample specific metadata). A further complication is that any analysis is not static but an iterative process. We therefore deemed it necessary to anchor a robust database at the centre of our software. In CytoPy, *Projects* are instantiated and housed within this database, which serves as a single dynamic data repository that is then accessed continuously throughout the subsequent analysis. For the architecture of this database, we chose a document-orientated database, MongoDB [20], where data are stored in JavaScript Object Notation (JSON)-like documents in a tree structure. Document-based databases carry many advantages, including simplified design, dynamic structure (*i.e*. database fields are not ‘fixed’ and therefore resistant to unforeseen future requirements) and easy to scale horizontally, thereby improving integration into web applications and collaboration. In this respect, CytoPy depends upon MongoDB being deployed either locally or via a cloud service, and MongoEngine [21], a Document-Object Mapper based on the PyMongo driver.

### 2.2 Framework overview

An overview of the CytoPy framework is given in Figure 1 including a recommended pathway for analysis, although individual elements of CytoPy can be used independently. CytoPy follows an object-orientated design with a document-object mapper for both commitment to, and collection from, the underlying database. The user interacts with the database using an interface of several CytoPy classes, each designed for one or more tasks. CytoPy is algorithm agnostic, meaning new autonomous gating, supervised classification, clustering or dimensionality reduction algorithms can be introduced to this infrastructure and applied to cytometric data using one of the appropriate classes. CytoPy makes extensive use of the Scikit-Learn and SciPy [22] ecosystems. Throughout an analysis, whenever single cell data are retrieved from the database, they are stored in memory as Pandas DataFrames that are accessible for custom scripting at any stage.

**Figure 1.**
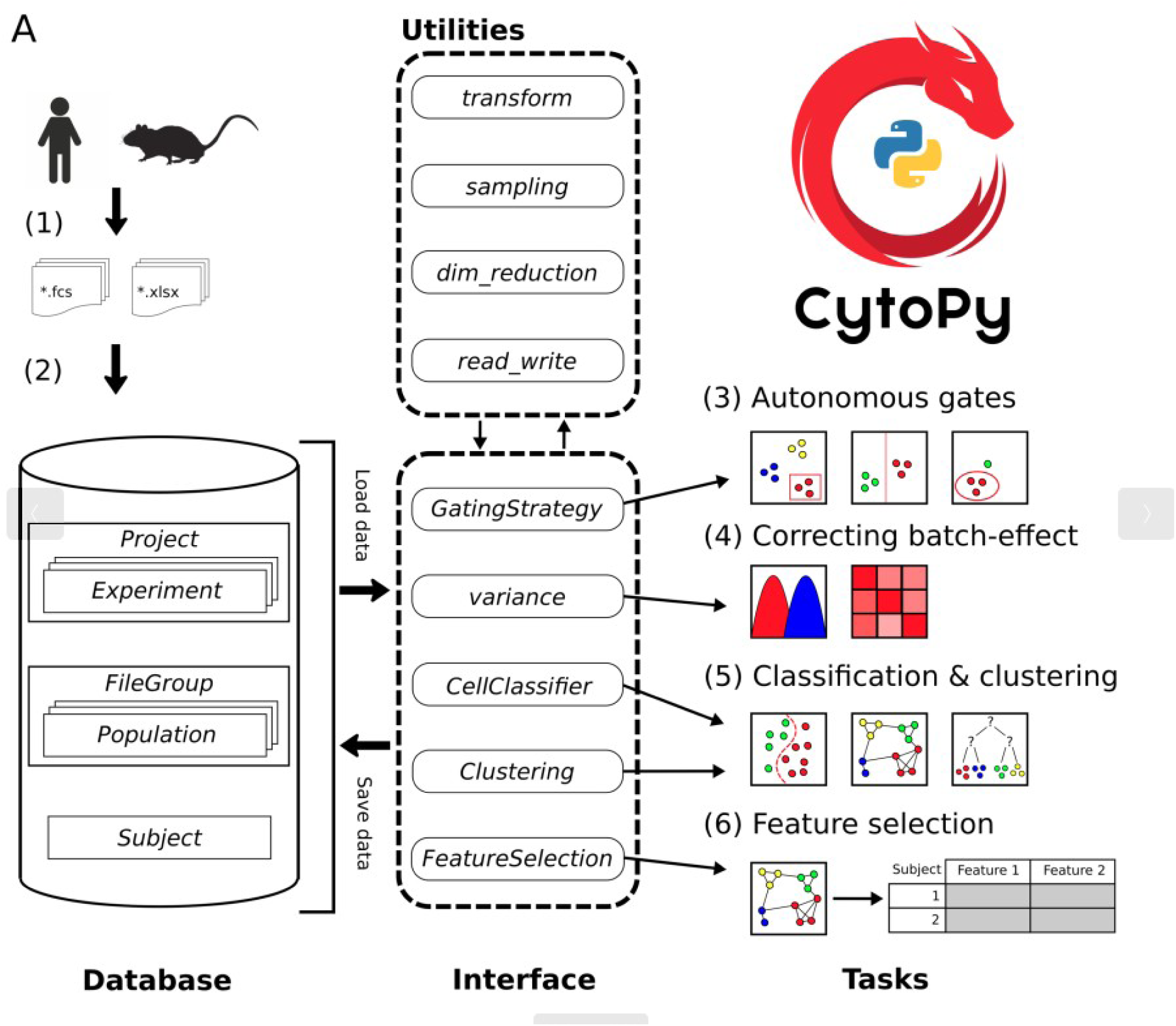
Overview of the CytoPy framework. Single cell data and experiment/clinical metadata (1) are used to populate a project within the CytoPy database (2). The CytoPy database models analytical data in MonogDB documents (cylinder), and an interface of CytoPy classes retrieves and commits data to this database (dotted rounded rectangle). Utility modules perform regular tasks such as data transformations and sampling throughout the framework. The components of this interface can be used independently, but the recommended workflow is as follows: (3) autonomous gates identify a ‘clean’ population of interest from where to start analysis, (4) batch effect is visualised, quantified, and corrected using the Harmony algorithm, (5) supervised and unsupervised algorithms classify cells into groups of similar phenotype, and finally (6) a feature space of cell population descriptive statistics is generated and feature extraction/selection methods deployed to identify a predictive signature that characterises an endpoint of interest.

Following the steps in Figure 1, a typical analysis in CytoPy would be performed as follows (functions show in italics and class names are shown in italics and title-case):

(1) Single cell data are generated and exported from the flow cytometer; CytoPy supports FCS (Flow Cytometry Standard) files version 2.0, 3.0 and 3.1, but additionally supports the introduction of data using a Pandas DataFrame object, therefore supporting wider formats, although this requires that the end user generates this object with suitable formatting. Experimental and clinical metadata are collected in tabular format either as Microsoft Excel document or Comma Separated Values (CSV) files, with the only requirement being that metadata be in ‘tidy’ format.
(2) A *Project* is defined and populated with the single cell data and accompanying metadata. A *Project* contains one or more *Experiment* documents, each defining a set of staining conditions. Each subject (*e.g*. a patient, a cell line, or an animal) has a *Subject* document containing metadata that are dynamic and have no restriction on the data stored within, and that are associated to one or several *FCSGroup* documents. Each *FCSGroup* document contains one or more FCS files (or *DataFrames*) associated to a single biological sample collected from the subject. This document contains all single cell data, ‘gated’ populations, clusters and meta-information that attains to a single ‘sample’, which also includes any isotype or Fluorescence-Minus-One (FMO) staining controls. Compensation is applied to single cell data at the point of entry using either an embedded spillover matrix or a provided CSV file. It should be noted that data are stored on a linear scale with a variety of transformations available during subsequent analysis; this provides flexibility in analysis as the user can compare the effects of different transformations, including the commonly used biexponential and hyperbolic arcsine transformations (compensation and transformations are implemented with the FlowUtils package [23]).
(3) Any cytometry analysis will require that single cell data be cleaned of debris and artefacts. We recommend FlowAI [24] be used independently of CytoPy prior to analysis to improve the quality of data. Within CytoPy, manual or autonomous gates can be employed to identify cell populations in two-dimensional space, replicating traditional manual analysis conducted with tools such as FlowJo. We recommend autonomous gates be used for eliminating doublets, dead cells, and debris, and to select a starting population for analysis; for instance, in a mixture of immune cells this could be the T cell population (CD3^+^ live single lymphocytes). Autonomous gates are applied with the *GatingStrategy* module and cell populations are then stored within the database as *Population* documents embedded within a *FileGroup.* These *Population* documents record the index of events belonging to a population, detail how they were identified, and the conditions in which they were identified such as transformations applied to linear space, *e.g.* biexponential transformation of axis.
(4) Batch effects are common and must always be addressed prior to analysis. If the batch effect is minimal the investigator can consider pooling data and modelling the distribution of single cell data directly. If batch effects are considerable, the investigator should include methods to alleviate this prior to further analysis. The *Variance* module of CytoPy provides methods to visualise and quantify batch effect. Autonomous gates provide methods to address batch effect through landmark registration, but technical variation can be addressed at a global level using the Harmony algorithm [25], implemented in CytoPy using the harmonypy package [26].
(5) Multiple strategies can be employed to classify cells based on a common phenotype. Strategies such as autonomous gating and supervised classification are biased by the training data provided (and the gating strategy used to label those data) whereas high-dimensional clustering is an unsupervised method that groups cell populations according to their phenotype. CytoPy offers both supervised classification through the *CellClassifier* class and high dimensional clustering through the *Clustering* class, so that variables can be generated from either or both strategies. These classes provide objects that are algorithm agnostic, allowing for the introduction of any function with specific signatures, whilst also providing much convenient functionality for visualising and critiquing results; this includes but is not limited to, cross-validation, learning curves, heatmaps, plotting with dimension reduction, and common metrics. Importantly, the results of either strategy generate common *Population* documents that are committed to the database and can then be used as input to any additional analysis or visualisations.
(6) Once cells have been classified, the user can test their hypothesis. The single cell data are summarised into a ‘feature space’, summary statistics that describe the cell populations. This generates a large number of variables, many of which will be either uninformative or redundant. Filter and wrapper methods are available through the *feature_selection* module finding only those variables that are important for predicting a biological or experimental endpoint. This module deploys methods from the discipline of interpretable machine learning, from simple L1-regularised linear models and decision trees, to complex modelling and interpretation through permutation feature importance and SHapley Additive exPlanations (SHAP) [27]

## 3. Results

### 3.1 Identifying significant batch effect in blood T cell subsets

To validate and exhibit the individual elements of CytoPy we decided to use the framework to identify T cells subsets in PBMCs isolated from whole blood. 14 individuals were chosen (based on availability of data) from a local study of patients undergoing peritoneal dialysis, 4 of whom presented with symptoms of acute peritonitis whereas the remainder were stable and asymptomatic (see supplementary methods). The objective was to identify T cells (single live CD3^+^ cells) in the first instance and then subsequently identify CD4^+^ T helper cells, CD8^+^ cytotoxic T cells, Vα7.2^+^ CD161^+^ mucosal-associated invariant T (MAIT) and Vδ2^+^ γδ T cell subsets. These populations were chosen to test a range of functionality: the ability to identify large and easy to distinguish cell types (CD4^+^ and CD8^+^ T cells), and more complex cell populations that can be rare in some patients and difficult to identify reliably in two-dimensional space (Vδ2^+^ γδ T cells and MAIT cells). Performance was compared to manual gates decided by user expertise.

The chosen study obtained patient material over 24 months resulting in significant batch effect. Supplementary Figure S1 shows the variation between individual fluorochromes, exhibiting ‘drift’ in the fluorescent intensity of multiple channels. Figure 2 shows UMAP plots of data from individual patients compared to a reference patient (blue); to choose a reference the pairwise Euclidean distance of a set of covariance matrices for each sample was computed and the sample with the smallest average distance to every other sample was chosen [9]. The UMAP plots revealed common structures shared between patients but a lack of alignment, suggesting the infiltration of noise from technical variation.

**Figure 2.**
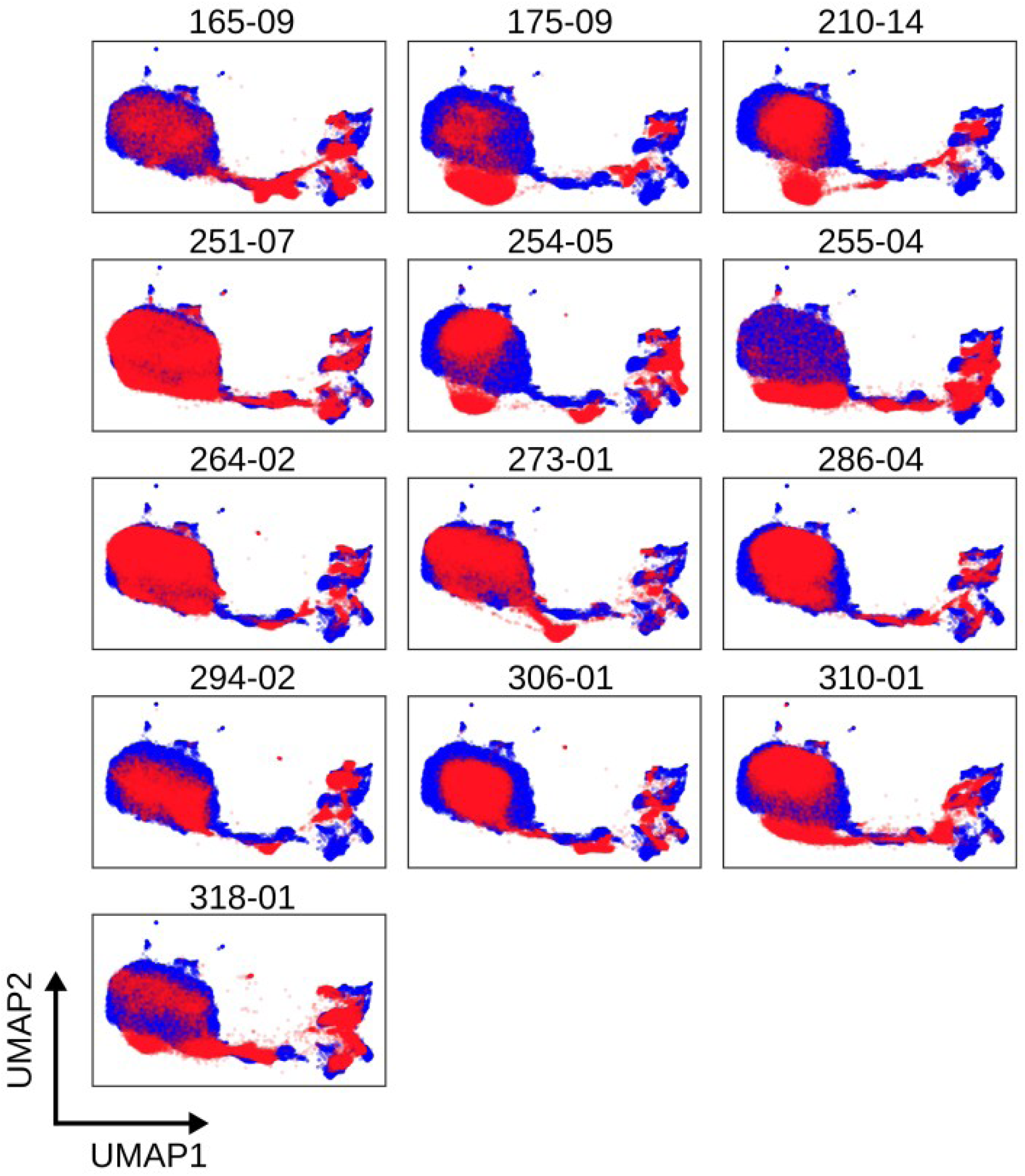
UMAP plots revealing batch effect in T cell staining of whole blood. A reference sample (blue) is chosen as the ‘average’ sample in Euclidean space. A low dimension embedding of this sample is made using UMAP (other algorithms are available in CytoPy, *e.g*. PCA, PHATE, tSNE) and samples for comparison are projected into this same space (red), demonstrating ‘drift’ in cell populations between patient samples. Each plot depicts results obtained with cells from an individual patient; numbers shown are unique patient sample identifiers.

### 3.2 Autonomous gates reliably identify T cell subsets despite batch effect

CytoPy replicates traditional manual gating using autonomous gates building on previous examples in the literature [1], [2]. The *Gate* object is used to implement a single algorithm for the identification of one or more cell populations in one or two dimensions. *Gate* objects can then be ‘stacked’ within a *GatingStrategy*, saved to the database and applied in sequence to subsequent data. Each *Gate* is defined using some example data and an algorithm chosen that best encapsulates the population of interest. The example data that a *Gate* is defined on acts as a reference to the expected populations in subsequent data. On exposure to new data, the algorithm is reapplied and the resulting populations matched the expected populations from the example data. Multiple algorithms are available for autonomous gates and are discussed in detail in the supplementary methods.

A challenge when defining autonomous gates is the choice of hyperparameters that will generalise beyond the chosen example data; this is further exacerbated by batch effects. CytoPy employs two techniques to overcome this issue: hyperparameter search and landmark registration. Hyperparameter search allows the user to specify a range of hyperparameters to use when a *Gate* is applied to new data. An exhaustive search is performed across all permutations of chosen hyperparameters resulting in a set of populations. For populations defined by a polygon gate, the convex hull of each population is computed and the population with the minimum Hausdorff distance to the population in the example data is chosen. For populations defined by a positive/negative threshold in one or two dimensions, the median fluorescent intensity of each population is computed and the population with the minimum Euclidean distance to the original example population is chosen. This is repeated for each population captured by a *Gate*.

Additionally, a user can apply landmark registration at the point of application of a *Gate*. First described in the context of cytometry data by Hahne *et al*. [28], landmark registration is used to align data to a reference by finding a warping function that aligns landmarks in their estimated probability density functions (Supplementary Figure S2). Landmarks are identified as points of maximum density and grouped by a K means algorithm [28]. In CytoPy we follow the method described by Finak *et al.* [29] and perform local normalisation when a *Gate* is applied.

Whilst accounting for batch effect with hyperparameter search and landmark registration, we applied autonomous gates to identifying T cell subsets in PBMCs (Supplementary Figure S3). Figure 3 shows a comparison of the number of events identified by autonomous gates (x-axis) compared to the same population identified by manual gates (y-axis), where each data point is an individual patient. Autonomous gates showed good conformity with manual gates, even for small and difficult to distinguish Vδ2^+^ γδ T cells and MAIT cells.

**Figure 3.**
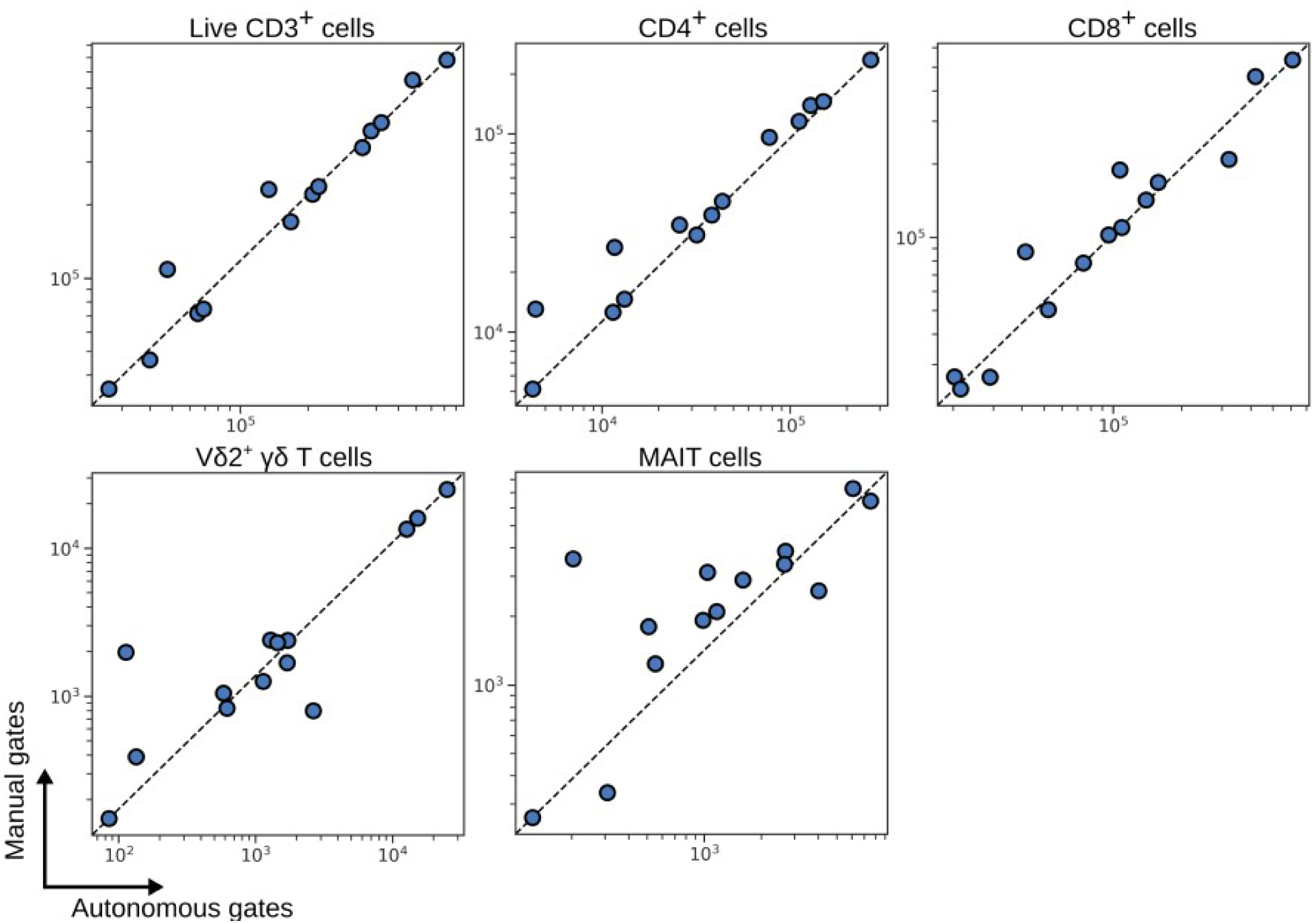
Number of events captured by autonomous gates for blood T cell subsets compared to the same subsets as defined by manual expert gates. Each symbol depicts results obtained with cells from an individual patient.

### 3.3 Batch effect can be addressed ‘globally’ using the Harmony algorithm

Despite the success of autonomous gates for the identification of T cell subsets in the wake of significant batch effect, they are heavily biased by the choice of example data when defining *Gate* objects and by the choice of reference for landmark registration. An alternative approach to addressing batch effect is to try to align cell populations between individual subjects in high dimensional space prior to analysis. There have been several methods proposed with this objective [30], most prominently applied to single cell RNA sequencing data, although some examples such as SAUCIE [31] demonstrate application to cytometry data.

We decided to implement the Harmony algorithm [25] given its ability to scale to large data and its transparent hyperparameters. Harmony was originally described as being applied to low-dimensional embeddings. This is necessary for RNA sequence data where the number of available features can be in the thousands or tens of thousands but is not necessary for cytometry data with only a dozen or more parameters. Therefore, we exposed the original data to the Harmony algorithm, after removal of debris, doublets, and dead cells. Biexponential transformation followed by scaling of each parameter to unit variance (by subtracting the mean and dividing by standard deviation) was performed prior to batch effect correction.

The performance of Harmony when applied to our T cell population (as identified by autonomous gates) from PBMCs is shown in Figure 4. Harmony has a range of hyperparameters that influence its behaviour. We found that default values for more of these parameters provide good performance but σ should be varied to improve performance on cytometry data; this hyperparameter influences the entropy regularisation term of the soft-clustering step of the algorithm and as it approaches zero, clustering is more alike to hard K means clustering. We chose an optimal value of 0.2 for σ whilst limiting the number of iterations to 5. The quality of batch correction is assessed by observing the distribution of the local inverse Simpson’s Index (LISI) and a UMAP embedding of single cell data before and after running Harmony (Figure 4A); LISI is effectively the number of batches in a cell’s local neighbourhood [25]. The objective here was to redistribute LISI such that the local neighbourhood around a cell contains a greater representation of different batches, without over-correcting and distilling biological variation that differentiates groups of subjects.

**Figure 4.**
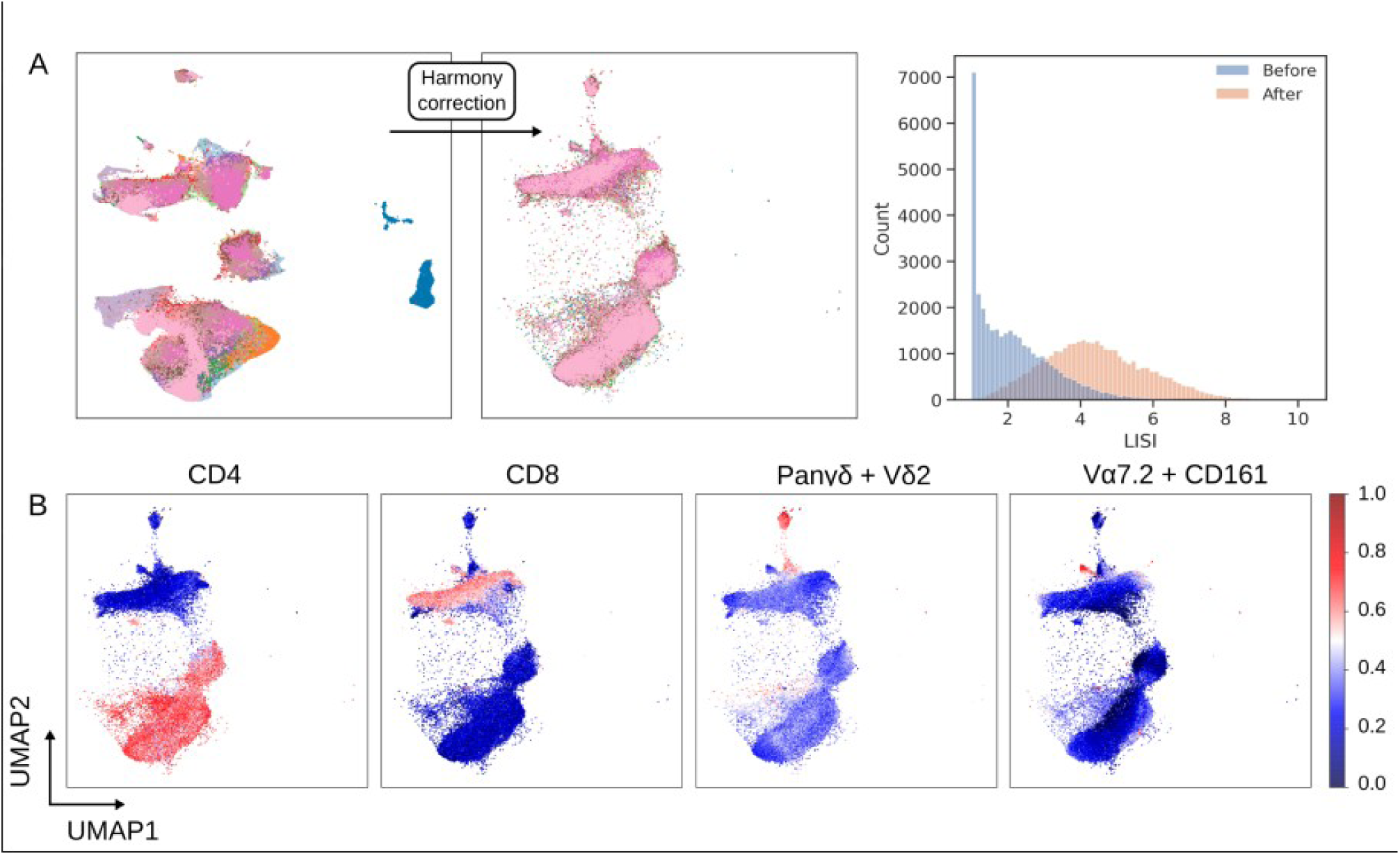
Batch correction using the Harmony algorithm. (A) Single cell UMAP plots are coloured by cell origin, where each colour represents a unique patient. Shift in batch membership in the local neighbourhood of cells is shown by the change in the UMAP plot after Harmony is applied and by the shift in LISI distribution. (B) Cell population structure is conserved after correction as shown by the shape of latent variables UMAP1 and UMAP2, and the distribution of the cell surface markers CD4, CD8, the linear combination of Panγδ and Vδ2 (to identify Vδ2^+^ γδ T cells), and the linear combination of CD161 and Vɑ7.2 (to identify MAIT cells).

The UMAP plots in Figure 4A show that prior to applying Harmony large communities of cells consist of single batches whereas after application these communities are diffused yet maintain a topology of separate cell populations. The concern with batch correction is over-correction that disrupts the biological meaning of the single cell data, but Figure 4B and 4C demonstrates that biological meaning is conserved, with distinct cell populations identified by their structure and lineage markers.

### 3.4 Supervised methods can replicate the performance of autonomous gates

Considering hundreds of thousands of data points can be obtained for each subject, cytometry data lends itself well to a supervised classification approach for identifying cell populations. Supervised classification of cell populations is exposed in CytoPy through the *SklearnCellClassifier* and *KerasCellClassifier* classes, which inherit from the *CellClassifier* class. Objects of this class can accept any classifier that conforms to/supports the Scikit-Learn API (such as XGBoost) or a Keras [17] model. Many convenient methods are pre-built into these objects and predictions can be saved as *Population* objects, providing compatibility with all other tools in the CytoPy framework.

To benchmark this supervised approach, we compared four native classifiers from Scikit-Learn (logistic regression, linear discriminate analysis, support vector machine with radial kernel, and K-nearest neighbours), XGBoost [32], and a deep feed-forward neural network built with Keras to classifiers reported from the Flow Cytometry: Critical Assessment of Population Identification Methods competition (FlowCAP) [33]. Algorithms were chosen from a range of classifier families based on their popularity in the literature. Supplementary Table S1 reports the weighted F1 score for each classifier across the five example datasets from FlowCAP. A deep neural network, with the architecture described by Huamin *et al.* [9], showed good performance as previously reported. XGBoost additionally showed exceptional performance, which again highlights the ability of this classifier to generalise to a wide range of use-cases.

The FlowCAP competition provides example data that have been heavily pre-processed and is not representative of data encountered in large clinical studies. We therefore decided to test the utility of XGBoost on the classification of T cell subsets. Since batch effect has been accounted for using Harmony, we pooled data from all available samples to generate training data that were manually labelled using the gating infrastructure within CytoPy. Tools for assessing the performance of a classifier such as cross-validation, learning curves, and confusion matrices are provided in CytoPy as convenient methods in *CellClassifier* (Supplementary Figure S4). As illustrated in Figure 5, XGBoost is capable of identifying T cell subsets and is comparable to manual gating. Since batch effect correction with Harmony involves a down-sampling step, comparisons are shown as the percentage of T cells as observed by manual gates vs populations identified by XGBoost.

**Figure 5.**
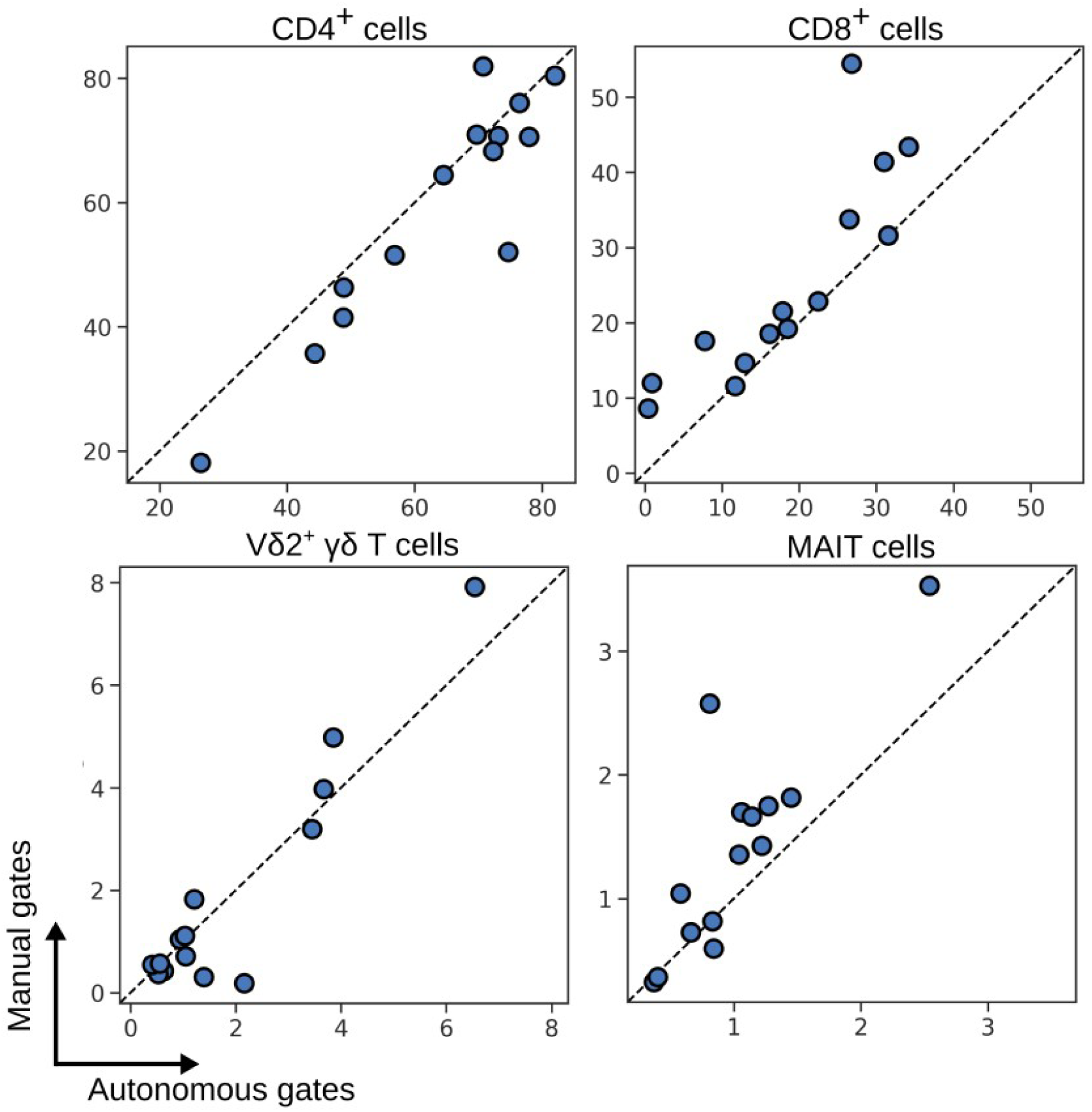
Percentage of blood T cell subsets as identified by XGBoost compared to the same subsets as identified by expert manual gates. Each symbol depicts results obtained with cells from an individual patient.

### 3.5 FlowSOM and Phenograph clustering for identifying T cell subsets after removal of batch effects

Autonomous gates and supervised classification are capable of identifying known populations of interest but are biased by the investigator’s understanding and expectations of the immune landscape. To diminish this bias, CytoPy encourages the use of unsupervised techniques alongside directed analysis.

Unsupervised clustering is a popular approach to identifying structures in single cell data, with techniques such as FlowSOM and Phenograph growing in popularity in the field of cytometry data analysis. Both algorithms are available in CytoPy, along with clustering algorithms from the Scikit-Learn ecosystem and consensus clustering. Any clustering algorithm can be applied within the framework to generate *Population* objects using the *Clustering* class and a function with a common signature and expected output. This design was chosen to future-proof CytoPy against further developments in the field so that new techniques will be easy to integrate using a simple wrapper function.

Clustering of the batch effect corrected T cells was performed using both FlowSOM and Phenograph by directing the *Experiment* towards a *Clustering* object and supplying the relevant functions. Each sample within an *Experiment* was clustered independently and then inter-sample comparisons were made through meta-clustering, as originally described by Levine *et al.* [4]. The results of meta-clustering are displayed in Figure 6 and demonstrate the ability of these algorithms to discern individual cell populations. The UMAP plots show each individual cluster as obtained from individual subjects but plotted in the same two-dimensional space and coloured by meta-cluster membership; the size of the data point corresponds to the proportion of events as a percentage of T cells in each individual. Most clusters are represented by a mixture of all subjects in an *Experiment* (Supplementary Figure S5) yet rare cell populations are under-represented in FlowSOM clustering; for instance, five patients had MAIT cells absent in clustering results despite being identified by manual gates (Figure 7). A comparison of the proportion of cells obtained by FlowSOM and Phenograph to the same cell type identified by manual gates showed that Phenograph gives preferable performance over FlowSOM. Despite this, Phenograph overestimated the proportion of Vδ2^+^ γδ T cells in a number of patients. This highlights the importance of using multiple techniques of both supervised and unsupervised classification when investigating cytometry data, and CytoPy simplifies this process.

**Figure 6.**
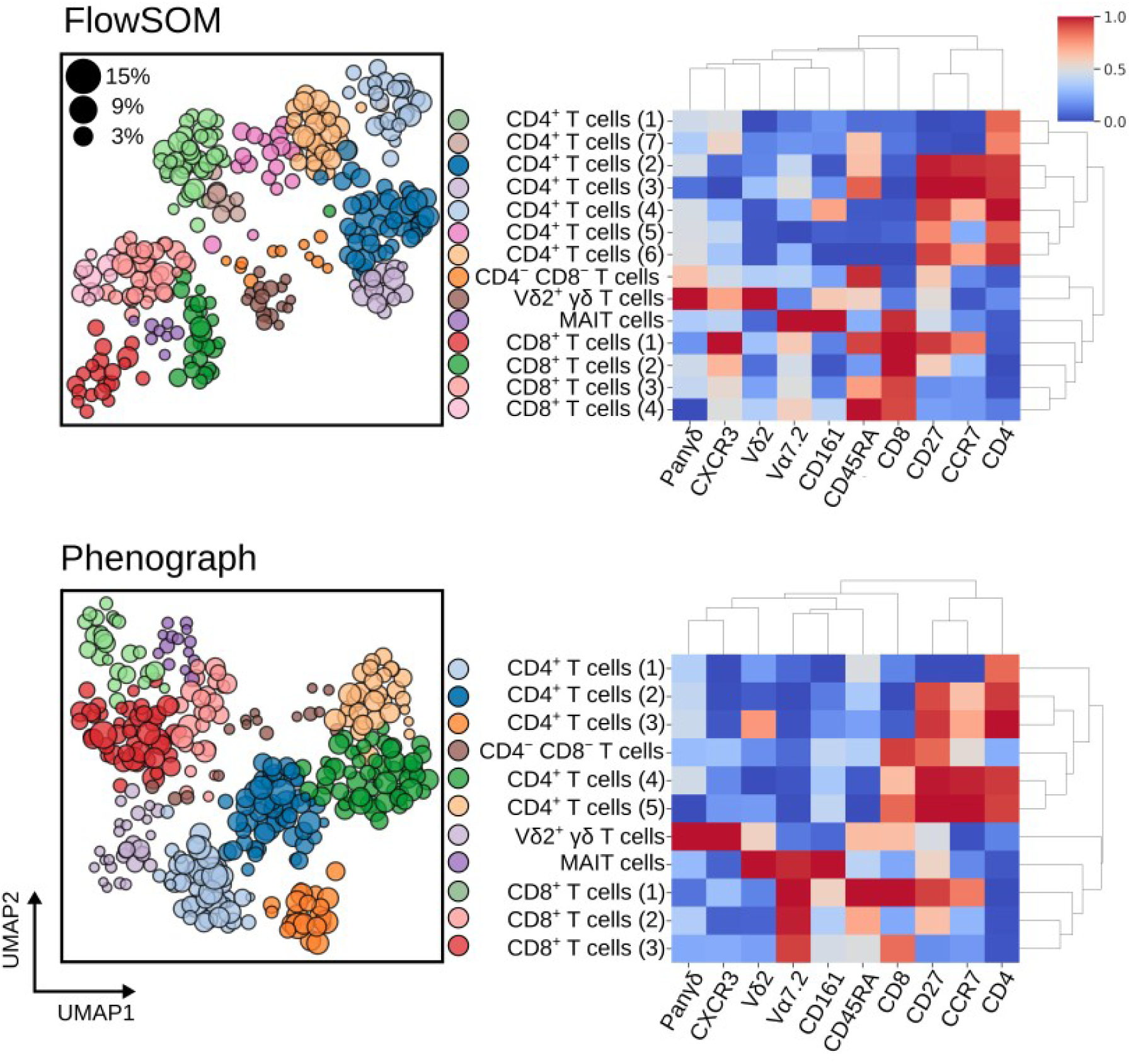
Meta-clustering results for FlowSOM (top) and Phenograph (bottom) when applied to blood T cells after batch effect correction with Harmony. Heatmaps show the normalised expression of cell surface markers for meta-clusters (clustered centroids of individually clustered patient samples). In the neighbouring UMAP plots, clusters from all patients are shown in the same embedded space and coloured by their meta-cluster membership. The size of each data point corresponds to the percentage of T cells this cluster represents in the patient it was derived from.

**Figure 7.**
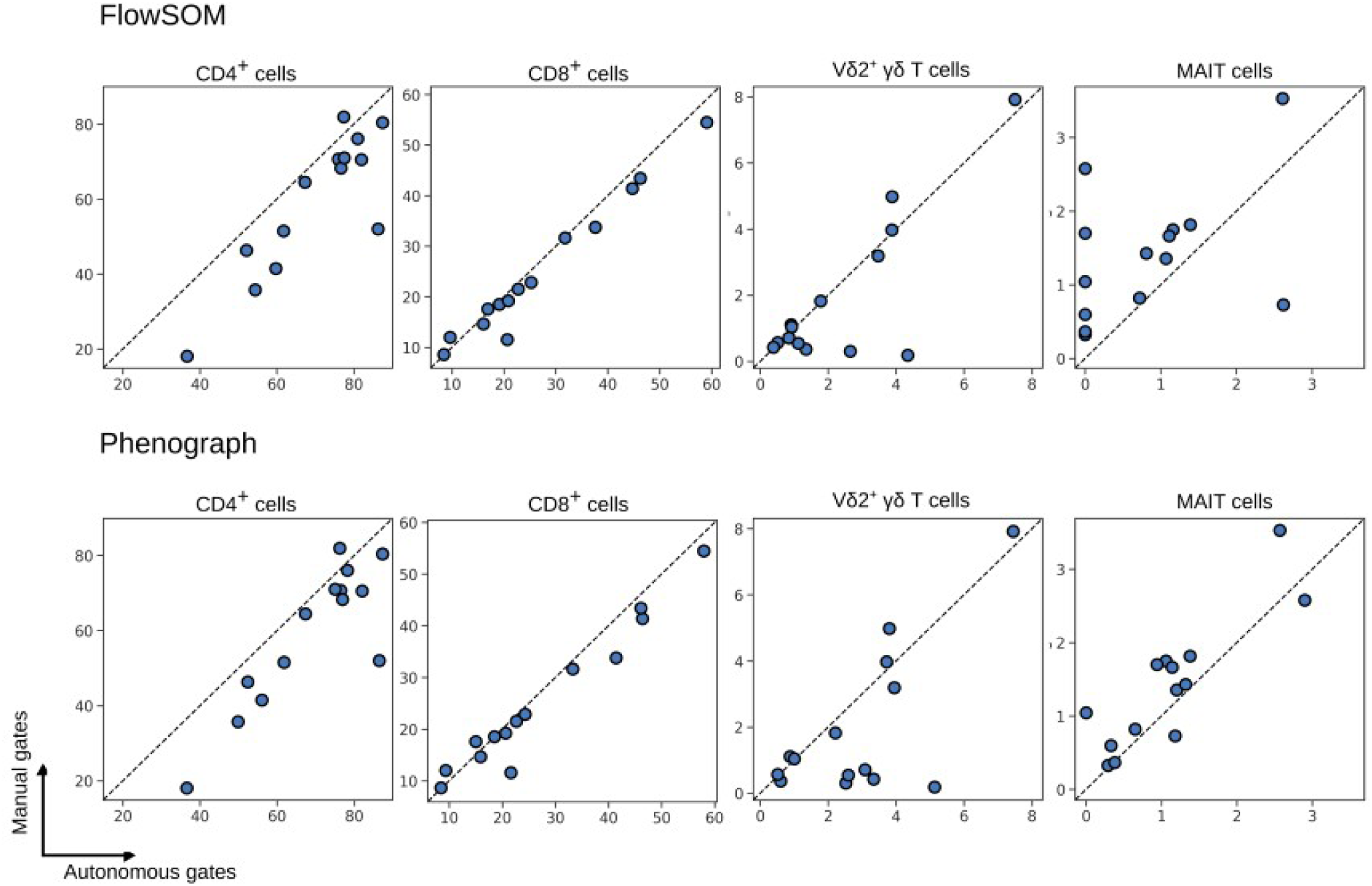
Percentage of T cell subsets as identified by FlowSOM (top) and Phenograph clustering (bottom), compared to the same subsets as identified by expert manual gates. Each symbol depicts results obtained with cells from an individual patient.

### 3.6 Implementing the CytoPy framework to identify an immune signature that differentiates patients with acute peritonitis from stable controls

To demonstrate the application of the entire CytoPy framework to an immunophenotyping project, we investigated the peritoneal effluent of patients undergoing peritoneal dialysis, some of whom presented with symptoms of acute peritonitis, with the objective to distinguish patients with acute peritonitis from stable controls based on their peritoneal immune signatures. This was chosen based on our long-standing expertise and published findings demonstrating the significance of the local immune response in recognising pathogen-specific patterns of infection [34] and the correlation between changes in myeloid populations and treatment failure [35].

T cells from PBMCs and the CD45^+^ fraction of cells from total effluent were obtained by autonomous gates prior to batch correction with Harmony (Supplementary Figure S6). XGBoost classification identified cell subsets using manual gates as training data displaying significant differences in the proportion of neutrophils and monocytes between patients with acute peritonitis compared to controls (Figure 8B). This was clarified in Phenograph and FlowSOM clustering and is exemplified in meta clustering UMAP plots (Figure 8B). The proportion of T cell subsets was not significantly different between stable controls and those presenting with acute peritonitis (Supplementary Figure S7).

**Figure 8.**
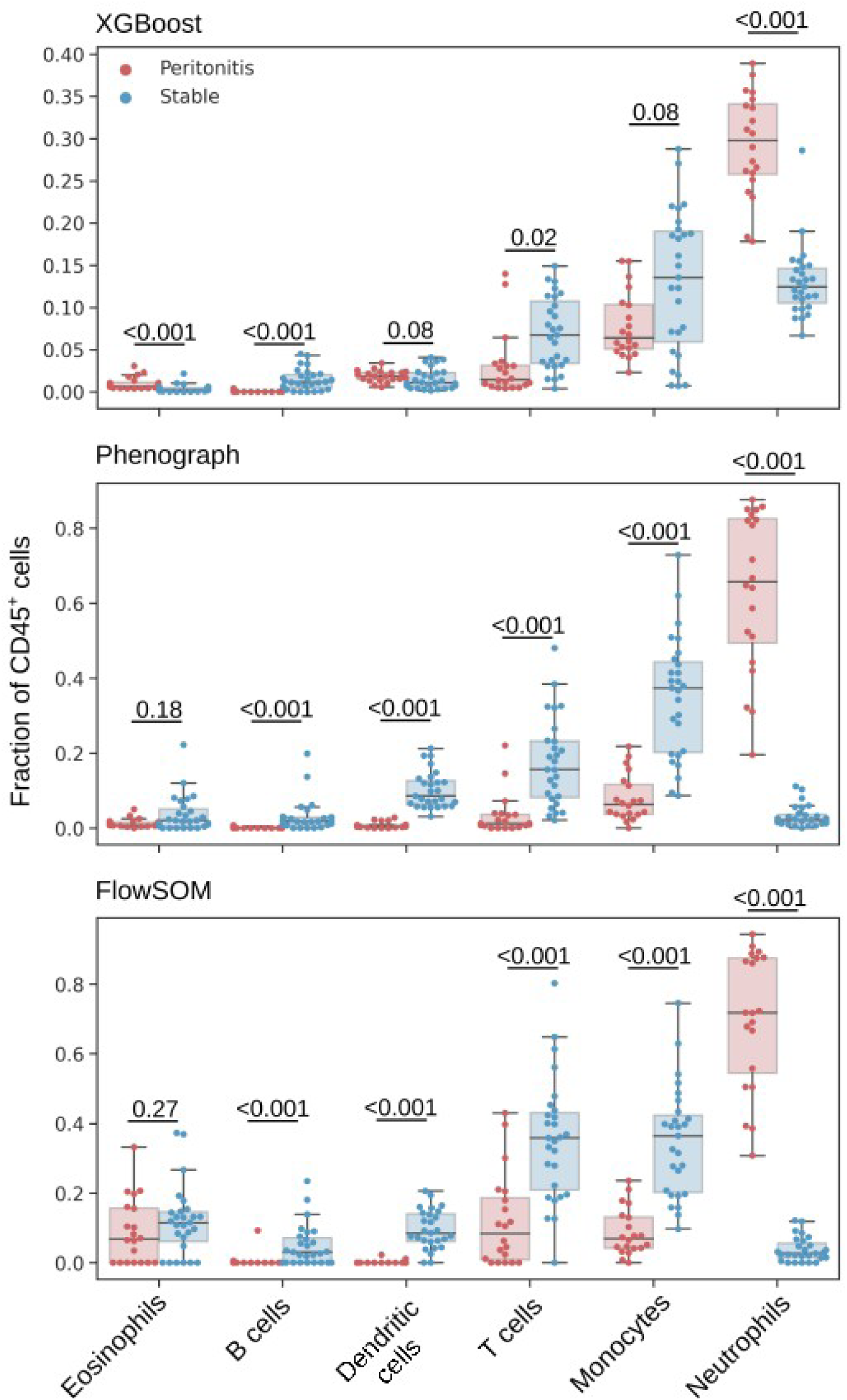
Leukocyte subsets as a fraction of CD45^+^ cells as identified by an XGBoost classifier (top), Phenograph clustering (centre) and FlowSOM clustering (bottom). Mann-Whitney U test were applied for comparisons between patients with acute peritonitis and stable controls, and p-values are reported after correction for multiple comparisons using Holm’s method (significance level was set as 0.05).

The ratio of cell populations observed by XGBoost classification and Phenograph, and FlowSOM clustering, conformed with one another (Figure 8). The average live CD45^+^ fraction (for T cells, B cells, monocytes, neutrophils, eosinophils) and average live T cell fraction (for CD4^+^, CD8^+^, Vδ2^+^ γδ T cells, and MAIT cells) across the three classification methods were pooled using the *feature_selection* module to generate a feature space representative of the local immune profile of the peritoneum. Age and gender were included in this feature space as potential confounding variables. High collinearity was observed between the fraction of CD4^+^ and CD8^+^ T cells, monocytes and DCs, and T cells and B cells (Figure 9A). CD8^+^ T cells, DCs and B cells showed low variability and were therefore removed from analysis. With the remaining features principal component analysis (PCA) was performed showing that patients with acute peritonitis were highly discernible from stable controls along the axis of the first principal component (Figure 9B). The absolute value of the coefficients for this component showed that neutrophils contributed the most to the observed variation. To confirm these findings, we generated a linear support vector machine with an L1 regularisation term using the *L1Selection* class. The regularisation parameter, *C*, was varied and the coefficient of each feature plotted; as the value of *C* decreases a sparse model is encouraged, eliminating features that do not contribute to the prediction. Figure 9C demonstrates that the fraction of neutrophils is the only feature to persist in a constrained model. CytoPy’s *feature_selection* module contains interpretable models for classification and regression problems, and its *DecisionTree* class can be used to demonstrate how the fraction of neutrophils alone can classify acute peritonitis (Figure 9D).

**Figure 9.**
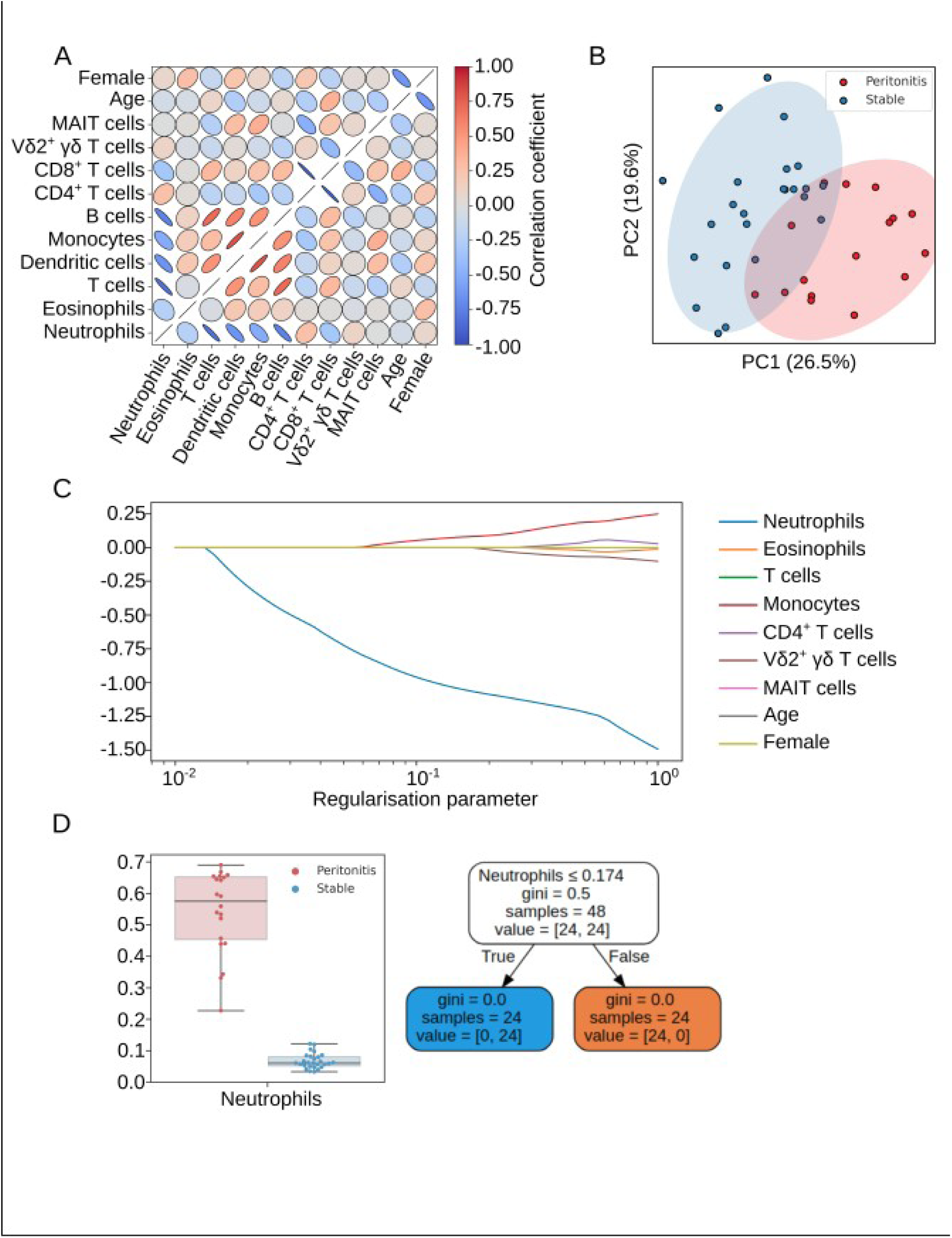
Feature selection process to reduce variables for predicting acute peritonitis. (A) Multicollinearity was addressed before generating linear models with redundant features removed prior to further analysis. (B) Principal component analysis shows that patients with acute peritonitis are discernible from stable controls. (C) L1 restricted modelling with a linear support vector machine reveals that neutrophils are the most predictive feature. (D) A simple neutrophils cut-off is predictive of acute peritonitis in this cohort and is demonstrated by a simple decision tree.

## 4. Availability and Future Directions

CytoPy represents a framework for the analysis of cytometry data that facilitates automated analysis whilst introducing robust data management and an iterative analytical environment. The present study shows the ability of CytoPy to characterise cell populations with high precision and the validation of the entire framework in identifying a known immune phenotype that distinguishes patients with acute peritonitis. This dataset was chosen based on our extensive experience with this sample type for over more than a decade. Initially acquiring such samples on a four colour BD FACSCalibur flow cytometer with two lasers and simple FSC/SSC settings [36], we later utilised an eight colour BD FACSCanto with three lasers and FSC/SSC area/height channels [37], and now in the present study took advantage of a 16 colour BD LSR Fortessa with four lasers and FSC/SSC area, height, width, and time [34], thus illustrating the technological advance in the field but also the increasing complexity of the data acquired. CytoPy exposes multiple techniques for the classification of cell populations in cytometry data with a simplistic design and a low-code interface. Autonomous gates provide a familiar interface with cytometry data whilst reducing the labour cost of analysis, despite this they’re biased by the investigator’s expectations of the data and computationally expensive. It is recommended that autonomous gates be employed for pre-processing and generating training data for supervised classifiers. Supervised classification offers a more efficient method for guided analysis, with training data provided in the form of a gated example. In contrast, unsupervised clustering is unbiased and offers exploratory analysis that can allude to the discovery of uncharacteristic cell populations or features that correlate with disease or experimental endpoints. In this study, we demonstrate that clustering algorithms such as FlowSOM and Phenograph were incapable of identifying rare cell populations for a small fraction of our cohort. This highlights the importance of not relying on a single method when engineering features from cytometry data. A cornerstone of CytoPy’s design is to expose multiple methodologies with minimal friction and provide consistent data structures to pool results. This strategy was employed for immune phenotyping peritoneal effluent and confirmed a striking increase in total neutrophils at the site of infection and a parallel decrease in the proportion of monocytes/macrophages, dendritic cells and T cells, in agreement with previous findings [34], [37], thereby validating the utility of CytoPy.

We have chosen to develop and maintain CytoPy in Python, a programming language with growing popularity in the bioscience domain. To date, Python has been lacking a framework for generalised cytometry data analysis offered by counterparts in R. CytoPy extends cytometry bioinformatics into the Python ecosystem by presenting an object-orientated infrastructure that is algorithm-agnostic and ready for deployment in the cloud. Compared to current solutions in R [2], [38], [39], CytoPy boasts a low-code interface and a data-centric design that enables rapid prototyping and comparison of analytical techniques, with seamless integration of metadata. Another popular solution for cytometry data analysis is CytoBank, which whilst supporting many popular algorithms and an accessible graphical user interface, is a propriety product that could limit uptake. In contrast, CytoPy is open-source and whilst offering popular algorithms, is also designed for expansion by the open-source community; new algorithms can be introduced with very simple wrapper functions to match existing signatures and expected data types.

The current version of CytoPy offers the most popular aspects of automated analysis of cytometry data, with autonomous gating, high-dimensional clustering and supervised learning, whilst also implementing Harmony [25] for batch effect correction. Future versions of CytoPy will expand on this to include algorithms such as SAUCIE [31] and BBKNN [40]. Another paradigm of immune phenotyping for predictive modelling is multiple-instance learning, whereby the single cell data from multiple patients are exposed to a model as a single matrix, with instances labelled by the kind of patient they originate from. This was successfully demonstrated by Hu *et al*. [13] to identify a signature predictive of latent cytomegalovirus (CMV) infection and by CellCNN [11] to identify paracrine signalling, AIDS onset, and rare CMV infection-associated cell subsets. It is our ambition to extend the capabilities of CytoPy to support this design.

As high-dimensional cytometry analysis continues to grow in popularity there is increasing demand for an analytical framework that is friendly for those who are new to programming, provides a database that directly relates experimental metadata to single cell data, and scales in a fashion that encourages collaboration and expansion. CytoPy meets all these criteria whilst remaining open-source and freely available on GitHub (https://github.com/burtonrj/CytoPy). Those wishing to collaborate with us or extend our software capabilities are invited to consult the documentation (https://cytopy.readthedocs.io/) and make a pull request on our GitHub repository.

## ACKNOWLEDGMENTS

We are grateful to all peritoneal dialysis patients for participating in this study, and to the clinicians and nurses for their cooperation. We also thank Chantal Colmont, Donald Fraser, Alexander Greenshields-Watson, Ann Kift-Morgan, Kristin Ladell, Oliwia Michalak and John Pulford for their help and advice. We would also like to acknowledge the open-source community that has contributed to the expansion of cytometry data analysis in Python, notably Peter Brodin, Brian Teague, Scott White, and Kamil Slowikowski. This research received support from the Wales Kidney Research Unit (WKRU), UK Clinical Research Network (UKCRN) Study Portfolio, Medical Research Council (MRC) grant MR/N023145/1, the Welsh European Funding Office’s Accelerate programme, and a School of Medicine PhD Studentship (to R.J.B.). The funders had no role in study design, data collection and analysis, decision to publish, or preparation of the manuscript.

## 5. Supplementary Methods

### 5.1 FlowCAP

Supervised classifiers in CytoPy were compared using data provided in the Flow Cytometry: Critical Assessment of Population Identification Methods (FlowCAP) challenge [33], where the challenge is to accurately separate cells into subsets based on single cell phenotype. The FlowCAP-I data consist of four human studies (graft-versus-host disease, diffuse large B-cell lymphoma, symptomatic West Nile virus infection, and healthy donors) and one mouse study (hematopoietic stem cell transplant). Data were labelled and pre-processing performed (removal of debris, dead material, and with fluorescence compensation applied) at source by the laboratory responsible for acquiring the original data. Here, classifiers were trained on 25% of data and classification performance tested on the remaining 75%. Performance was reported as the average of weighted F1 scores across all five datasets, where the F1 score for data with |*C*| set of possible classes is given as:

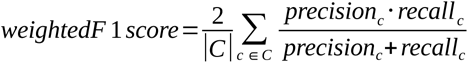

Six supervised machine learning algorithms, housed within CytoPy, were compared without hyperparameter tuning:

1. Logistic regression with balanced class-weights; implemented in Scikit-Learn version 0.24
2. Linear discriminant analysis without any shrinkage and number of components equal to either the number of classes or number of features, depending on which is minimum; implemented in Scikit-Learn version 0.24
3. Support vector machine with a radial basis function kernel without regularisation and γ as 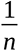 where *n* is the number of available features; implemented in Scikit-Learn version 0.24
4. K nearest neighbours classifier with *k* equal to 30; implemented in Scikit-Learn version 0.24
5. XGBoost using default parameters; implemented in xgboost version 1.2
6. Feed-forward neural network with three hidden layers of size 12, 6, and 3 nodes, L2 penalty of 1×10^−4^, softplus activation function on the hidden layers, softmax activation function of the outer most layer, and categorical cross-entropy as the loss function; implemented in Tensorflow Keras version 2.4

### 5.2 Patients

The study cohort comprised 21 adult individuals receiving peritoneal dialysis (PD) who were admitted between October 2016 and October 2018 to the University Hospital of Wales, Cardiff, on day 1 of acute peritonitis, before commencing antibiotic treatment (47.6% female; median age 53.0 years, range 30.0-86.0 years). 30 age and gender-matched individuals receiving PD and with no previous infections for at least 3 months served as stable, non-infected controls (53.3% female; median age 59.7 years, range 39.7-84.3 years). Subjects known to be positive for HIV or hepatitis C virus were excluded. Clinical diagnosis of acute peritonitis was based on the presence of abdominal pain and cloudy peritoneal effluent with >100 white blood cells/mm^3^. According to the microbiological analysis of the effluent by the routine Microbiology Laboratory, Public Health Wales, episodes of peritonitis were defined as infections caused by Gram-positive or Gram-negative organisms. Cases of fungal infection and negative or unclear culture results were excluded from this analysis. A summary of the bacterial culture results for patients with peritonitis are shown in Supplementary Table S2. All methods were carried out in accordance with relevant guidelines and regulations, and written informed consent was obtained from all subjects. Recruitment of PD patients was approved by the South East Wales Local Ethics Committee under reference number 04WSE04/27, and conducted according to the principles expressed in the Declaration of Helsinki. The study was registered on the UK Clinical Research Network Study Portfolio under reference numbers #11838 “Patient immune responses to infection in Peritoneal Dialysis” (PERIT-PD).

### 5.3 Flow cytometry

Peritoneal leukocytes were harvested from overnight dwell effluents and processed as described previously [34], [37]; samples were treated with DNase (Sigma; 1:2,500 dilution) when excessive debris was visually apparent. Leukocyte populations in total effluent were stained using monoclonal antibodies against CD1c, CD3, CD14, CD15, CD16, CD19, CD45, CD116, HLA-DR and Siglec-8 (Supplementary Table S3) and identified as CD45^+^ immune cells, CD3^+^ T cells, CD19^+^ B cells, CD15^−^CD14^+^ monocytes/macrophages, CD15^+^ neutrophils, CD15^−^CD14^+/−^CD1c^+^ dendritic cells, and CD15^−^SIGLEC-8^+^ eosinophils. T cell subsets in peripheral blood mononuclear cells (PBMCs) and in peritoneal effluent were stained after Ficoll (Ficoll-Paque PLUS; Fisher Scientific) separation of blood and peritoneal leukocytes, respectively, using monoclonal antibodies against CD3, CD4, CD8, CD161, TCR-Vα7.2, TCR-Vδ2, TCR-pan-γδ, CD45RA, CCR7 and CD27 (Supplementary Table S4). Cell acquisition by flow cytometry was performed using a 16 colour BD LSR Fortessa cell analyser (BD Biosciences). Live single cells were gated based on side and forward scatter area/height and live/dead staining (fixable Aqua; Invitrogen).

### 5.3 Autonomous gating

Autonomous gates inherit from the parent class *Gate* (providing access to common utilities such as data transformations) but are divided into the following classes to facilitate gating geometries:

1. The ***ThresholdGate*** divides data in one or two-dimensional space using a threshold of positivity (a straight line that divides data into positive and negative regions). Thresholds are found as regions of minimal density in the estimated probability density function of the observed data; estimated with a fast convolution-based kernel density estimation algorithm [41] with a Gaussian kernel and bandwidth estimated using the Silverman method.
2. The ***PolygonGate*** allows the user to apply any Scikit-Learn clustering algorithm to two-dimensional data; including the popular HDBSCAN algorithm [42]. Polygon gates are generated from the resulting clusters by computing their convex hull, the contents of this polygon are used to construct *Population* objects.
3. The ***EllipseGate*** allows the user to apply the probabilistic mixture model algorithms of the Scikit-Learn library to generate elliptical gates. For each component of the mixture model the covariance matrix is used to generate a confidence ellipse, surrounding data and emulating a gate. The ellipse is centred on the mean of the chosen component and oriented in the direction of the first eigenvector of the covariance matrix. The approximate likelihood of a data point falling within the bounds of the ellipse can be estimated using the chi-squared distribution. A hyperparameter, ‘conf’, is provided (default is 0.95) as the percentile of the chi-squared distribution to generate an ellipse where the length of the primary axis (the longest axis) is such that the chosen percentage of data attributed to this component is contained within the ellipse. This elliptical gate is then committed to the database as a polygon object.

### 5.4 Manual gating

T cells from whole blood were manually gated in FlowJo v10.7 (TreeStar) by two independent experts. The total number of events for each gate of interest were exported as a CSV file. The average number of events between the two independent analysts was used for comparison of automated methods to manual gating.

## Supplementary material

**Supplementary Figure S1.**
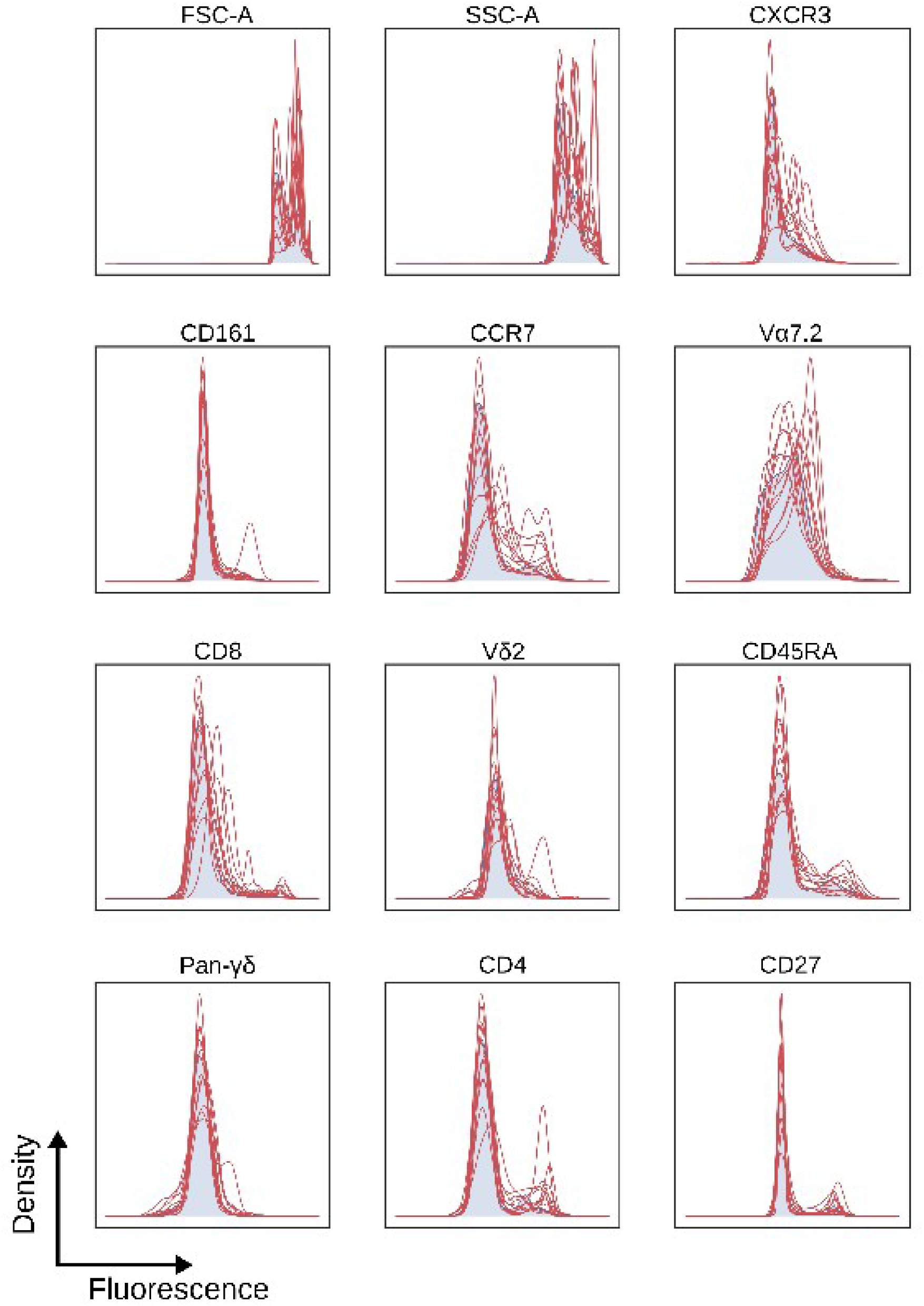
Individual fluorochrome inter-sample variation amongst PBMCs from 14 patient samples. Reference patient is shown in blue with subsequent patients overlaid as individual red lines.

**Supplementary Figure S2.**
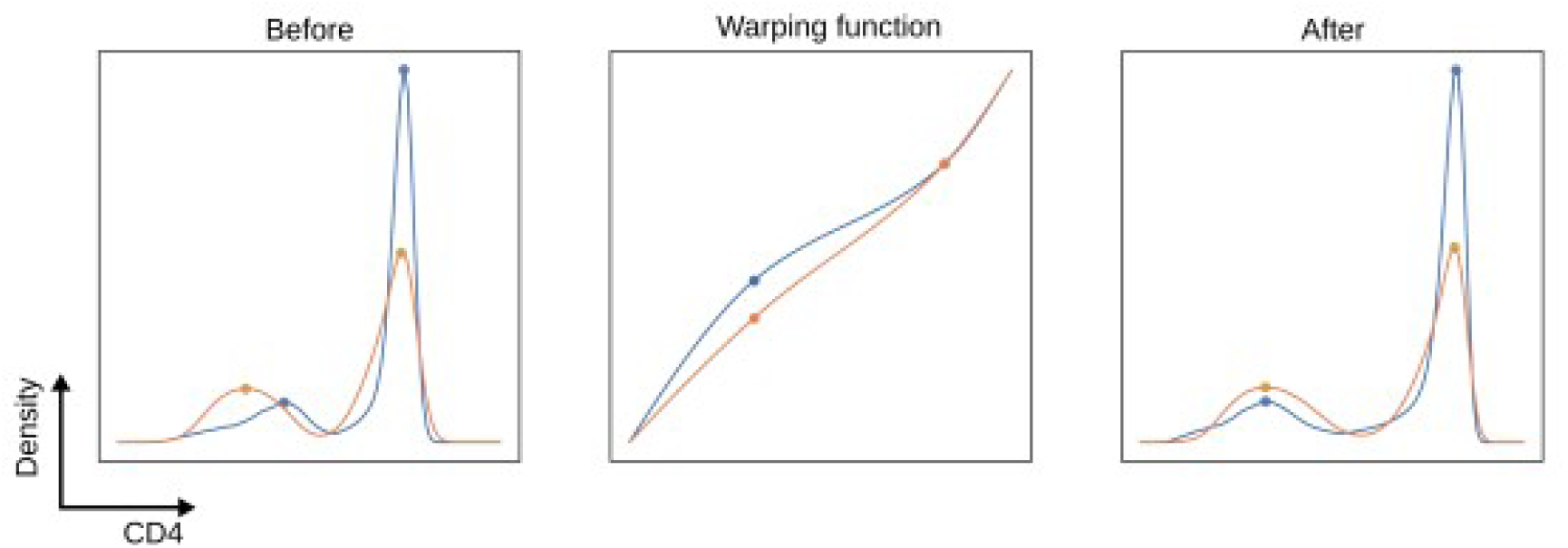
Example of landmark registration to align ‘peaks’ of high density in the probability density function (PDF) of CD4 expression on blood T cells for a target distribution and chosen reference. Left, target (orange) PDF compared to the reference (blue) prior to alignment. Centre, warping function defined between landmarks by taking a monotone cubic interpolation. Right, registered curve with aligned peaks obtained using function composition.

**Supplementary Figure S3.**
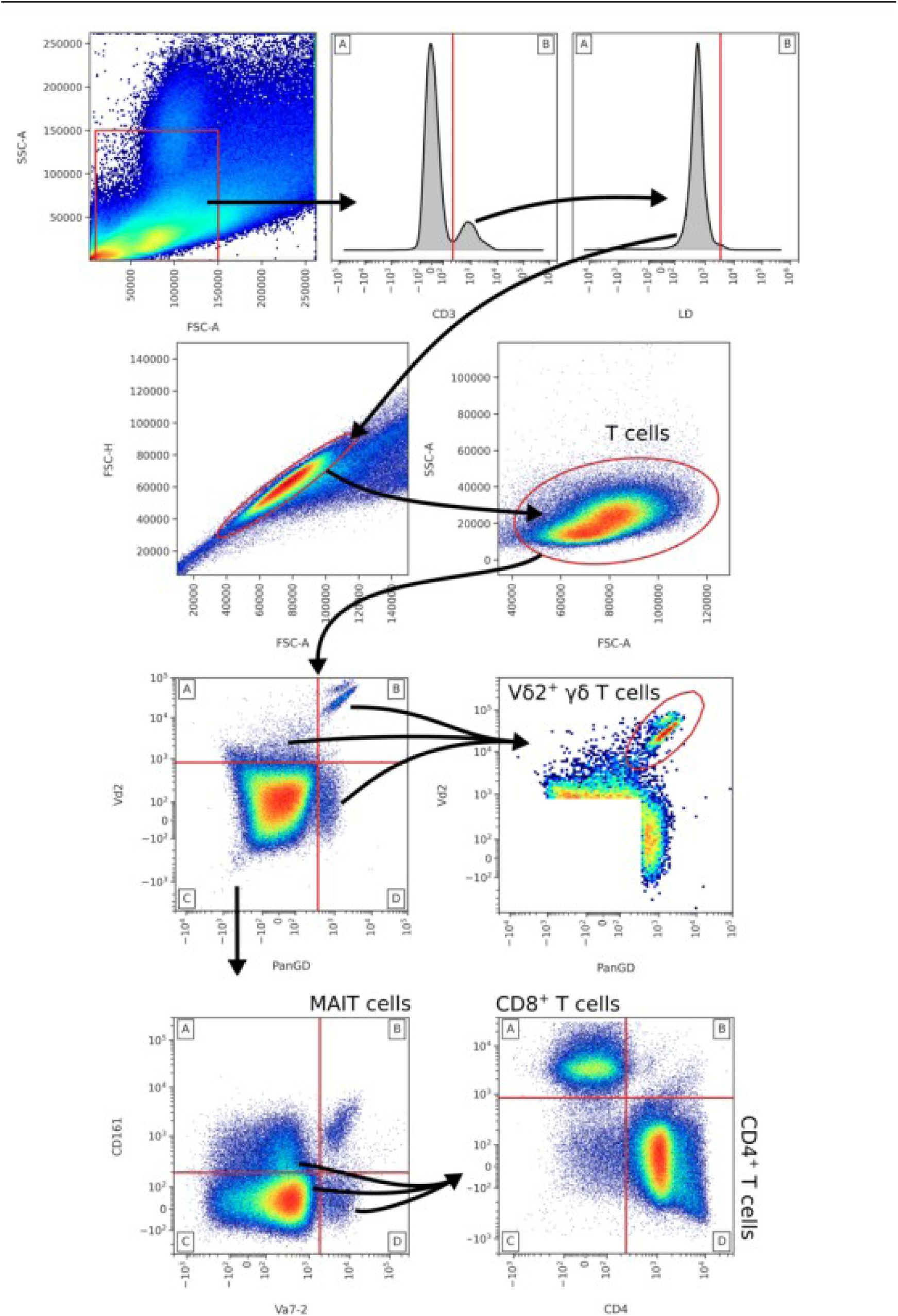
Autonomous gating strategy for identifying blood T cell subsets. Threshold (straight red line) gates are obtained using density-based algorithms as the *ThresholdGate* class, elliptical gates are obtained using the *EllipseGate* class and Gaussian mixture models with a varying number of components. Hyperparameter search and landmark registration was applied to all gates.

**Supplementary Figure S4.**
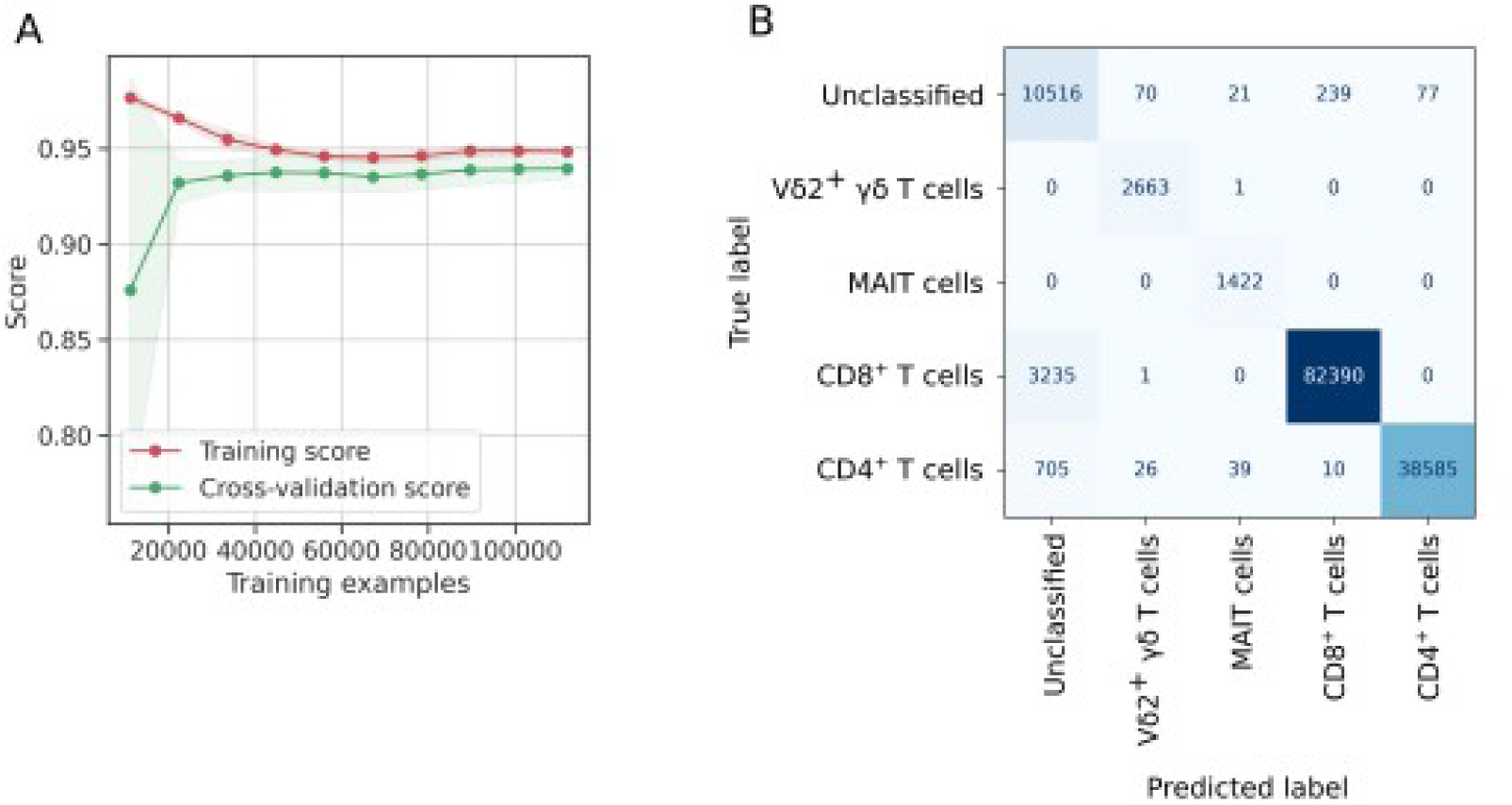
Example of a learning curve (A) for training XGBoost for identifying T cells subsets, and confusion matrix (B) for the same algorithm when exposed to validation data.

**Supplementary Figure S5.**
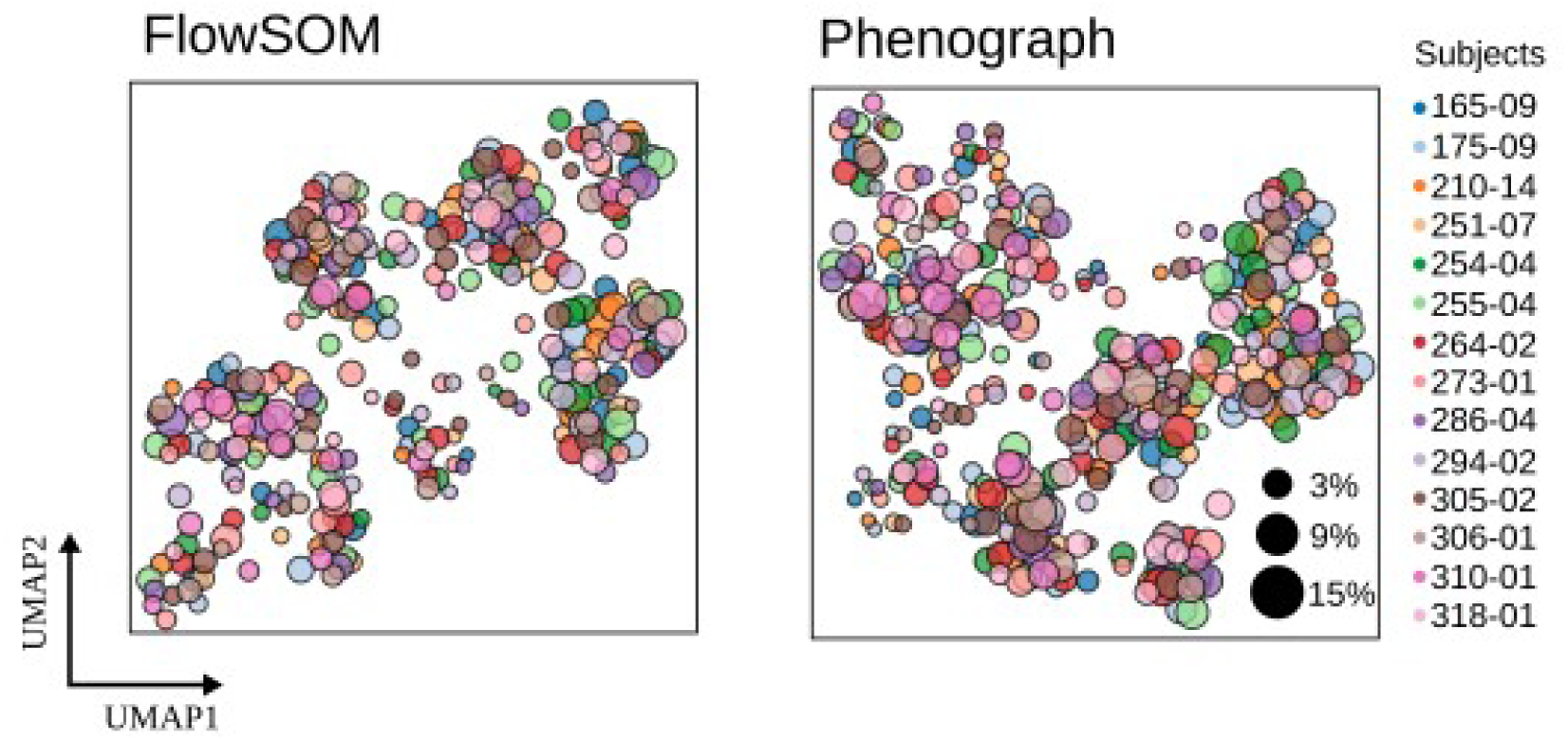
UMAP plots showing all clusters as obtained by FlowSOM (left) and Phenograph (right) and coloured according to patient origin. The size of the data points correspond to the % of T cells the given cluster represents from the respective patient. Subject numbers shown are unique patient sample identifiers.

**Supplementary Figure S6.**
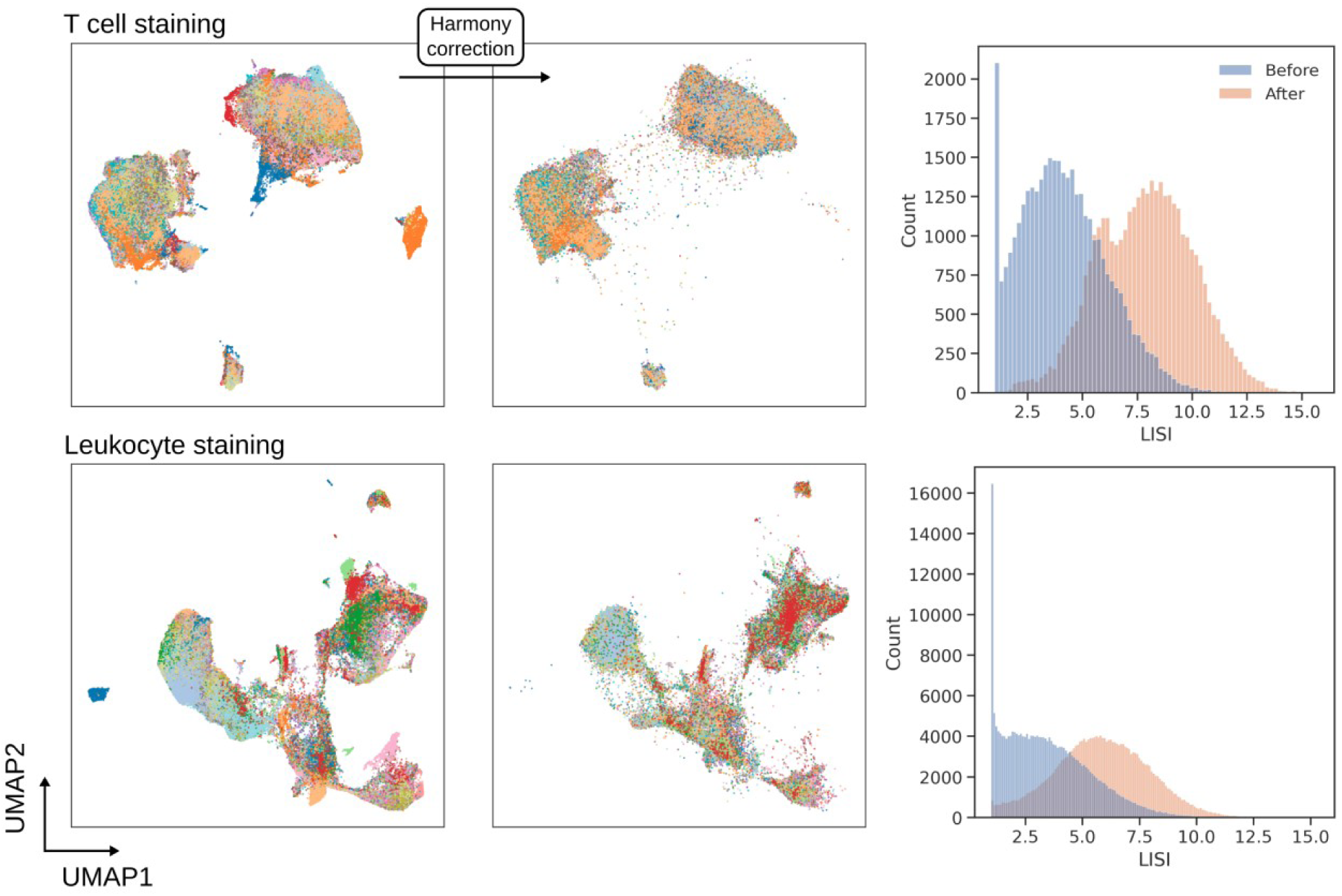
Batch effect correction with Harmony for peritoneal effluent stained for T cell subsets (top) and leukocyte subsets (bottom). Single cell UMAP plots are coloured to show the origin of cells where each colour is a unique patient. UMAP plots following correction and LISI distribution show the effectiveness of Harmony to correct for technical variation.

**Supplementary Figure S7.**
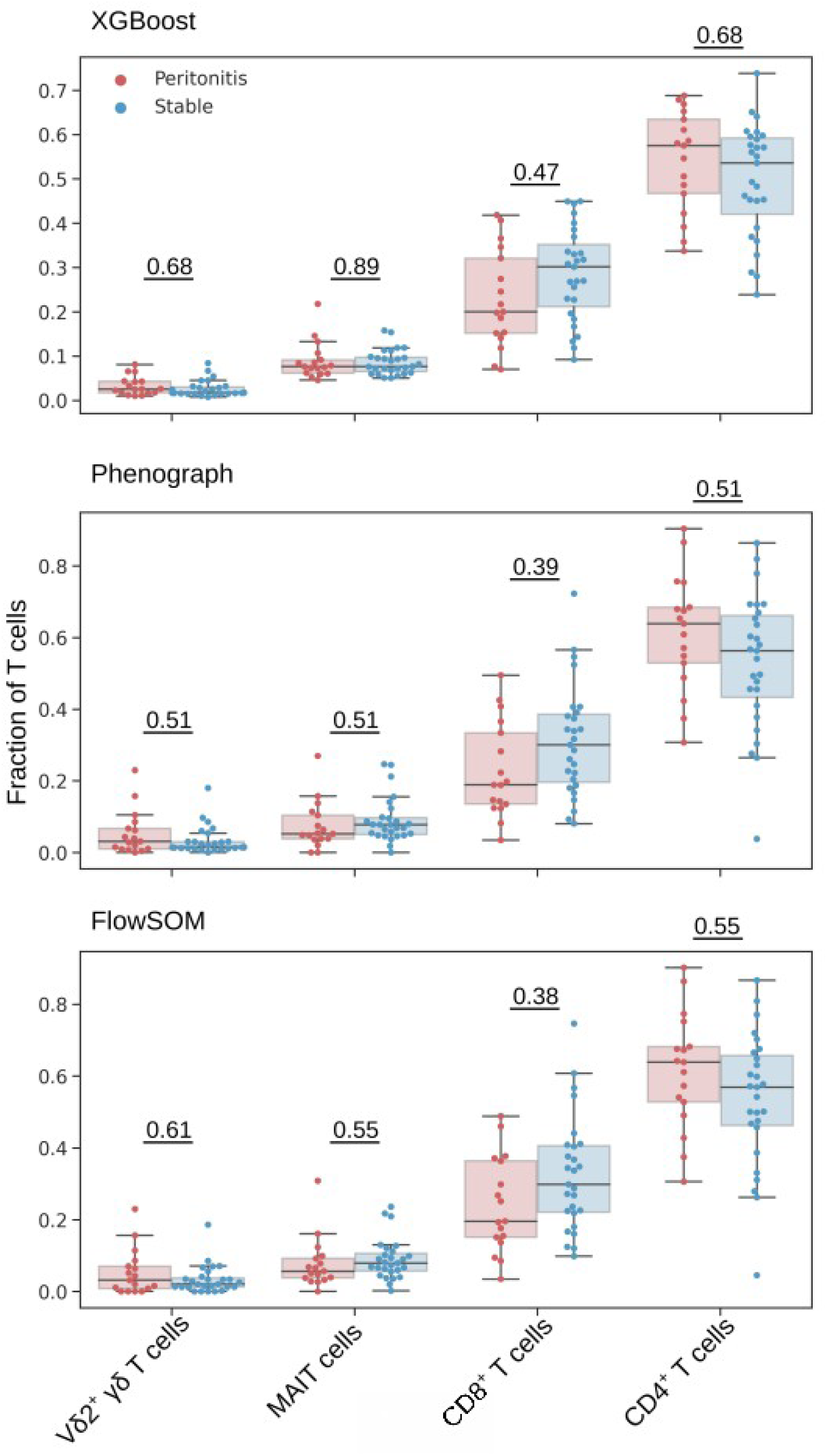
T cell subsets as a fraction of T cells, identified by an XGBoost classifier (top), Phenograph clustering (centre) and FlowSOM clustering (bottom). Mann-Whitney U tests were applied for comparisons between patients with acute peritonitis and stable controls and p-values are reported after correction for multiple comparisons using Holm’s method (significance level was set as 0.05).

**Supplementary Table S1.**
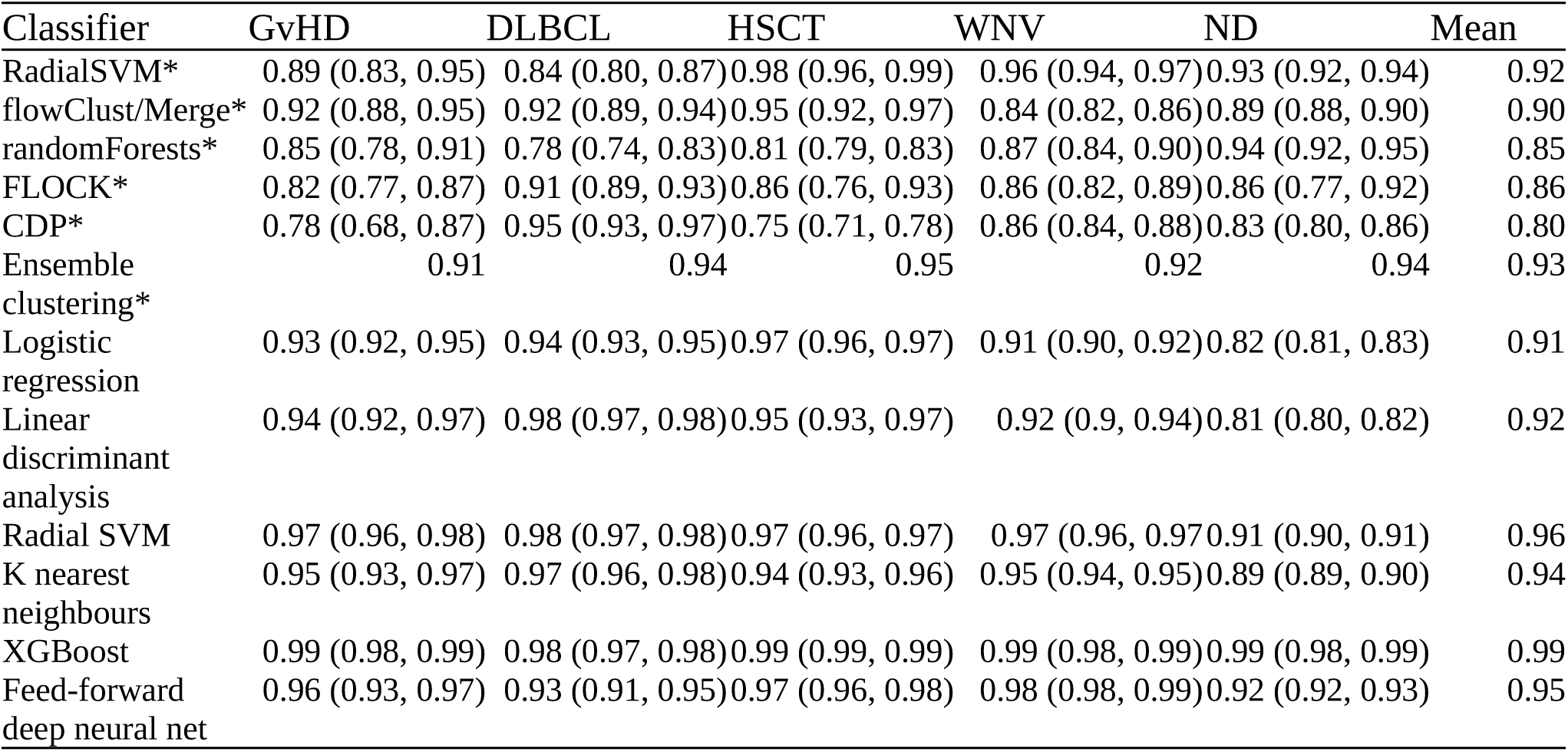
Performance of supervised classification algorithms for identifying cell populations from the FlowCAP competition data. *Performance from the original competition as reported by Aghaeepour N *et al.* [33]; all other algorithms are implemented through CytoPy.

**Supplementary Table S2.** Summary of microbiological culture results for peritoneal dialysis patients with acute peritonitis

| Culture result | *n* |
| --- | --- |
| Coagulase-negative *Staphylococcus* | 6 |
| Alpha-haemolytic *Streptococcus* | 3 |
| *Staphylococcus aureus* | 1 |
| *Escherichia coli* | 1 |
| *Streptococcus agalactiae* | 1 |
| *Corynebacterium amycolatum* | 1 |
| *Pseudomonas aeruginosa* | 1 |
| Yeast | 1 |
| Mixed growth | 2 |
| No growth/unknown | 4 |

**Supplementary Table S3.** Staining panel for T cells

| Marker | Fluorochrome | Manufacturer (Clone) |
| --- | --- | --- |
| CD3 | APC/Fire | BioLegend (UCHT1) |
| CD4 | PE-Cy5.5 | BioLegend (OKT4) |
| CD8 | Brilliant Violet 711 | BioLegend (RPA-T8) |
| CD161 | APC | Miltenyi Biotec (191B8) |
| Vα7.2 | Brilliant Violet 605 | Biolegend (3C10) |
| TCR-pan-γδ | PE-Cy5 | Beckman Coulter (IM2662) |
| Vδ2 | PE | BD Biosciences (B6 RUO) |
| CCR7 | Brilliant Violet 421 | BioLegend (G043H7) |
| CD27 | PE-Cy7 | BioLegend (M-T271) |
| CD45RA | PE Dazzle | BioLegend (HI100) |

**Supplementary Table S4.**
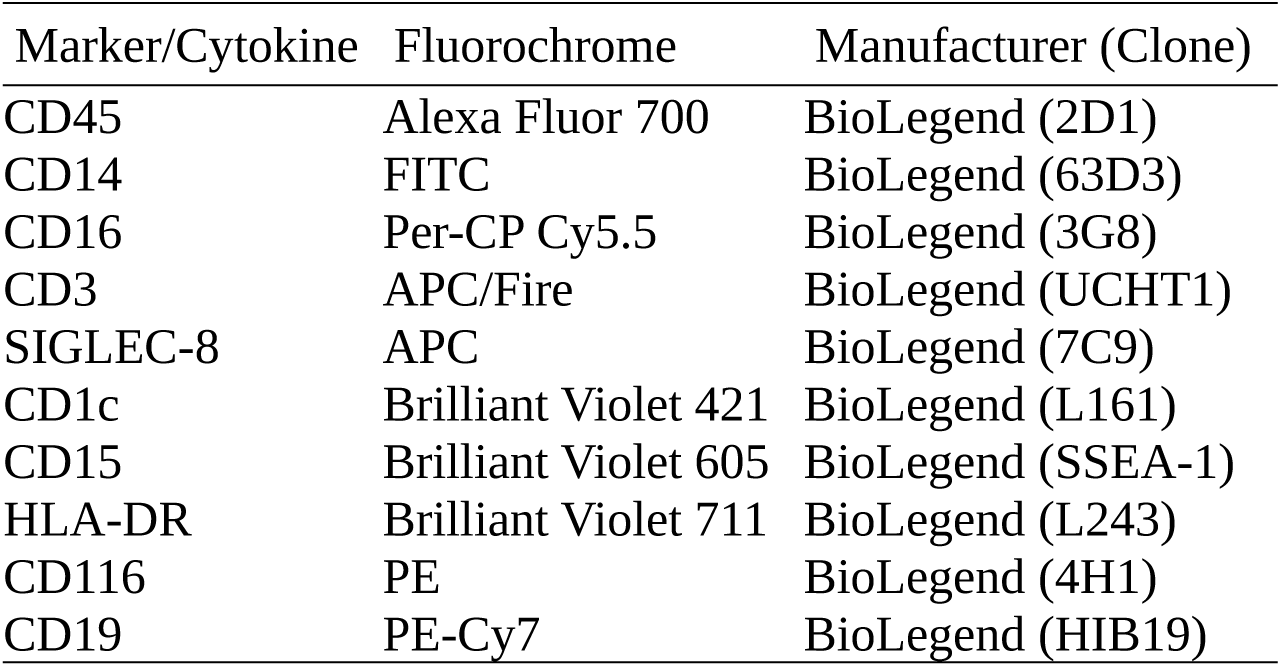
Staining panel for leukocytes

